# Mechanisms of adjustments to different types of uncertainty in the reward environment across mice and monkeys

**DOI:** 10.1101/2022.10.01.510477

**Authors:** Jae Hyung Woo, Claudia G. Aguirre, Bilal A. Bari, Ken-Ichiro Tsutsui, Fabian Grabenhorst, Jeremiah Y. Cohen, Wolfram Schultz, Alicia Izquierdo, Alireza Soltani

## Abstract

Despite being unpredictable and uncertain, reward environments often exhibit certain regularities, and animals navigating these environments try to detect and utilize such regularities to adapt their behavior. However, successful learning requires that animals also adjust to uncertainty associated with those regularities. Here, we analyzed choice data from two comparable dynamic foraging tasks in mice and monkeys to investigate mechanisms underlying adjustments to different types of uncertainty. In these tasks, animals selected between two choice options that delivered reward probabilistically, while baseline reward probabilities changed after a variable number (block) of trials without any cues to the animals. To measure adjustments in behavior, we applied multiple metrics based on information theory that quantify consistency in behavior, and fit choice data using reinforcement learning models. We found that in both species, learning and choice were affected by uncertainty about reward outcomes (in terms of determining the better option) and by expectation about when the environment may change. However, these effects were mediated through different mechanisms. First, more uncertainty about the better option resulted in slower learning and forgetting in mice, whereas it had no significant effect in monkeys. Second, expectation of block switches accompanied slower learning, faster forgetting, and increased stochasticity in choice in mice, whereas it only reduced learning rates in monkeys. Overall, while demonstrating the usefulness of entropy-based metrics in studying adaptive behavior, our study provides evidence for multiple types of adjustments in learning and choice behavior according to uncertainty in the reward environment.

## Introduction

Learning and choice in dynamic environments are shaped by a tradeoff between adaptability and precision (Farashahi et al., 2017a; Khorsand & Soltani, 2017; Soltani, Murray, et al., 2021). In stable environments, learning can be slow to allow more precise estimates of reward contingencies. In contrast, in unstable and thus less predictable environments, learning can become faster, slower, or show no change depending on where the decision maker falls in terms of adaptability and precision tradeoff. This is because faster learning enables faster updates but also comes at the cost of less precision, which may not be desirable in certain situations. Consistent with this, Behrens and colleagues (Behrens et al., 2007) have shown that more uncertainty in terms of volatility can result in higher learning rates, whereas others have found no evidence for such adjustments (Donahue & Lee, 2015, Farashahi et al., 2019). Instead, it was shown that volatility can decrease the weight of more uncertain information (estimated reward probability) relative to that of more certain information (reward magnitude) and improve encoding of task-relevant signals (Donahue & Lee, 2015; Massi et al., 2018; Farashahi et al., 2019).

Estimation and adjustments to uncertainty are tightly linked to detecting and learning regularities in the reward environment. Such regularities can modulate learning and decision making to enable the decision maker to anticipate and prepare for upcoming changes (Atilgan et al., 2022). Nevertheless, any regularity is accompanied by an inevitable uncertainty due to the stochastic nature of the reward environment. It is thus not surprising that the influences of different types of uncertainty on learning and the neural mechanisms by which these effects are mediated have been topics of much interest (Yu & Dayan, 2005; Behrens et al., 2007; Winstanley & Floresco, 2016; Faraji et al., 2018; Soltani & Izquierdo, 2019; Piray & Daw, 2021). For example, it has been shown that during a dynamic foraging task in mice, serotonin neurons in mice represent expected uncertainty and track unexpected uncertainty for updates (Grossman et al., 2022). In monkeys, neurons in the dorsolateral prefrontal cortex track the statistical variance of reward outcomes (risk) resulting from choices for specific options (Grabenhorst et al., 2019). Nonetheless, similarities and differences between adjustments to different types of uncertainty are not fully understood.

Understanding adjustments to uncertainty requires studying behavior in dynamic learning tasks that involve different types of uncertainty, for example the kind resulting from changing reward probabilities for different choice alternatives between blocks of trials. To reveal mechanisms underlying such adjustments, often choice data are fit using different reinforcement learning (RL) models that have been augmented with additional components to achieve better goodness-of-fit (Farashahi et al., 2017b, Wittmann et al., 2020; Womelsdorf et al., 2021; Grossman et al., 2022; Collins & Shenhav, 2022). Although RL models can reveal possible mechanisms underlying flexible behavior, fit of choice data requires many trials, which makes applying this method to study continuous adjustments in behavior very challenging.

To address this issue, we recently developed a set of new behavioral metrics based on information theory that can quantify consistency in trial-by-trial response to reward feedback using a relatively small number of trials (Trepka et al., 2021). These metrics provide a convenient, model-free tool to measure randomness in adaptive choice behavior and guide further model development. More importantly, these metrics can be computed from any segment of the task, enabling semi-continuous quantification of behavioral adjustments over time.

Here, we used these metrics to study adjustments in learning and choice behavior to different types of uncertainty in the reward environment. To that end, we re-analyzed data from mice and monkeys that performed similar dynamic foraging in which the selection of two choice alternatives was rewarded probabilistically, while baseline reward probabilities switched after a certain number of trials (referred to as the block length). We investigated the influence of uncertainty in determining the better option as well as uncertainty about the time of block switches on learning and choice using entropy-based metrics. These include overall entropy of choice strategy in terms of stay or switch, entropy of reward-dependent strategy, entropy of option-dependent strategy, and mutual information between reward outcome and choice strategy. While incorporating the probability of win-stay and lose-switch strategies, these metrics can better summarize local response to reward feedback, which then can be used to predict global choice behavior or its adjustments (Trepka et al., 2021).

With regard to uncertainty in determining the better option based on reward feedback, we hypothesized that increased uncertainty should result in slower learning to allow more accurate estimation of reward probabilities and/or, in more stochastic behavior, to enable more exploration. This could happen via different non-exclusive mechanisms such as a decrease in the learning rates and an overall increase in the stochasticity of choice. In addition, we hypothesized that when blocks are longer than expected, choice or adopted choice strategy would be more random, as animals, in the absence of any changes in the environment, more strongly expect changes in reward probabilities. Finally, expectation of block switches could interact and/or depend on the level of uncertainty in determining the better option.

Briefly, we found that stochasticity in choice behavior and response to reward feedback measured by entropy-based metrics were systematically affected by uncertainty about the better option and expectation of block switches in both mice and monkeys. Further, by fitting RL models to different groups of choice data identified by applying entropy-based metrics, we identified possible mechanisms by which different types of uncertainty influence behavior across the two species.

## Methods

### Experimental paradigms

To examine whether and how animals use regularities and uncertainty of the reward environment to guide their behavior, we reanalyzed data from mice and monkeys performing two dynamic foraging tasks with variable block lengths. There were key similarities in these experiments that allowed us to compare performance directly across species. First, while baseline reward probabilities remained fixed within a block of trials, probability of reward assignment on the unchosen option increased over time in a similar fashion in both tasks. Second, the block length changed from block to block but the range of block lengths were similar across the two species: with 40–100 trials in mice and 50–150 trials in monkeys.

#### Mouse experiment

In the mouse experiment (Bari et al., 2019), water-restricted mice were habituated for 1–2 days and head-restrained for the task. Eighteen male C57BL/6J (The Jackson Laboratory, 000664) mice aged 6–20 weeks were used in the experiment. A “go-cue” was signaled by an odor, after which the mice made a leftward or rightward lick toward one of the two water ports for a probabilistic reward (**Figure 1A**). Reward was a drop of water (2–4 μL). In 5% of trials, a “no-go cue” was signaled by a different odor, after which licks were neither rewarded nor punished. Each trial was followed by an inter-trial interval (ITI), of which the duration was drawn from an exponential distribution with a rate parameter of 0.3 with a maximum of 30s.

**Figure 1.**
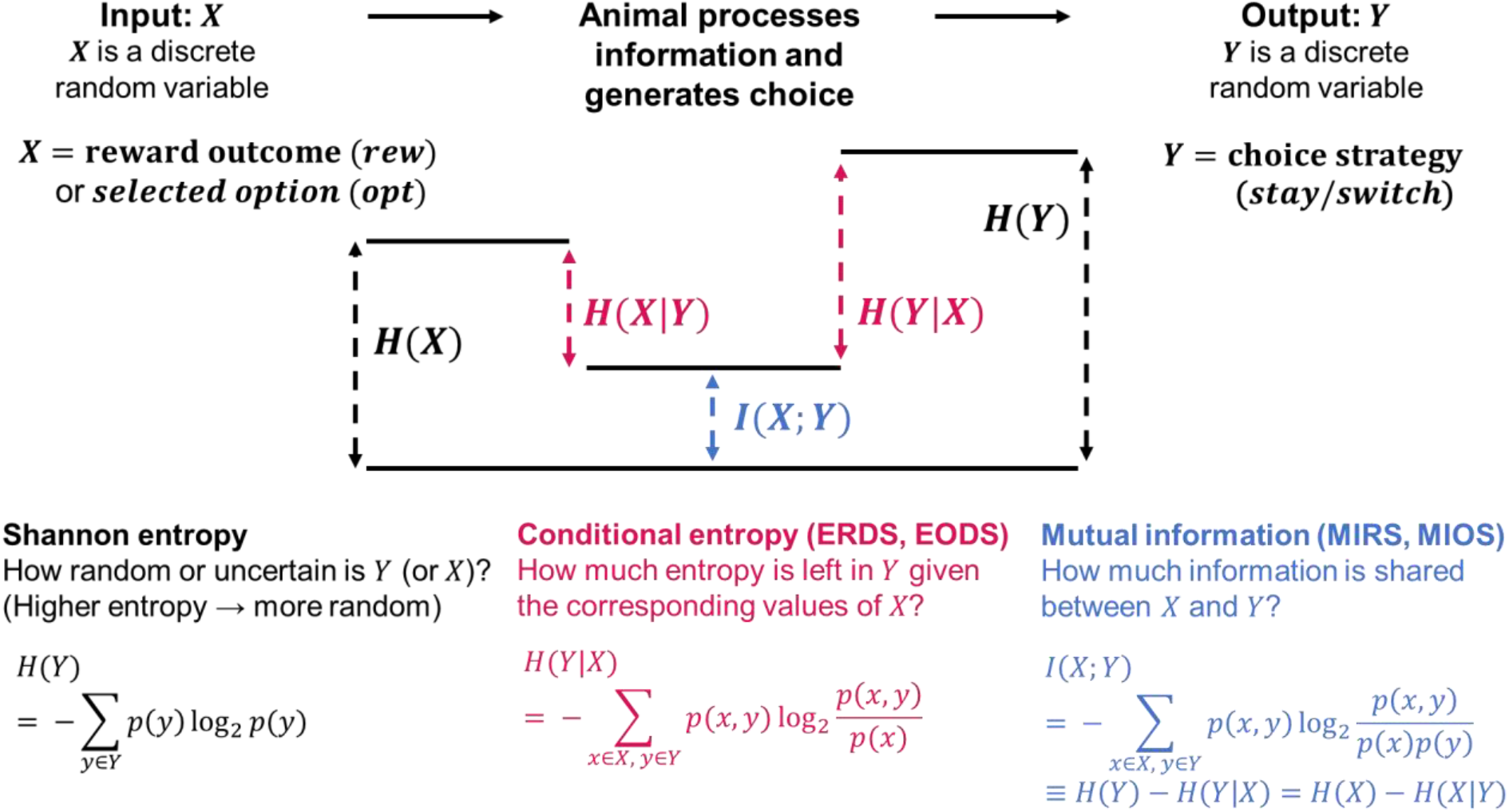
Application of information theory to studying choice behavior under uncertainty. The stochastic nature of the reward environment results in uncertainty in reward outcome that can be measured using Shannon entropy (*H*(X)). The animal used information in the environment to generate choice that can be quantified as repeating the previous choice (stay) or switching to the alternative choice option (for two-alternative choice tasks). Randomness in choice can also be measured using Shannon entropy (*H*(Y)). The link between previous reward outcome and animals’ strategy can be quantified using conditional entropy (e.g., *H*(Y|X)) measuring uncertainty in strategy given reward outcome in the previous trial, or mutual information (I(X; Y)) measuring shared information.

Reward probabilities were assigned to the left and right water ports, with one port yielding a higher reward probability (better option) than the other (worse option). Rewards were baited, such that if a reward was assigned to a particular side and was unchosen, the reward would remain there until that side was selected. This rule implies that the reward probability of an unchosen option becomes larger with the number of trials unchosen, mimicking ecological conditions in which unvisited foraging options will become more rewarding with time. Reward probabilities were chosen from 16 different sets of reward schedules (**Figure S1C**). The majority of sessions (88.8%) used two sets of reward probabilities equal to 0.4 and 0.1 (40/10) and 0.4 and 0.05 (40/5). The two reward probabilities were held constant for a fixed number of trials (a block) and reversed (block switches or reversal) without any cues to the animals. The lengths of each block were drawn from a uniform distribution spanning 40–100 trials. On rare occasions when mice demonstrated strong side biases, block lengths were shortened or lengthened.

For a given day of experiment, mice performed one of two versions of the task, a two-probability task (523 sessions) and a multiple-probability task (only 5 sessions). In the two-probability task, only two reward probabilities were assigned from the pool of possible probabilities and reversed for a given session. In the multiple-probability task, the better and worse options were still reversed after block switches, but more than two reward schedules were used for a given session. These probabilities were chosen such that the ratios would equal {8:1, 6:1, 3:1, 1:1}, to match parameters from the previous experiment by Sugrue and colleagues (Sugrue et al., 2004). A majority of the mice (16 out of 18) performed the two-probability task only. For the other two mice, multiple-probability sessions were randomly interleaved with two-probability sessions. In total, 18 mice performed 3768 blocks during 528 sessions (mean 7.13 block switches per session). Here, we focus our analyses on the two-probability task only because most of the mouse experiment was done using this task, and there was a significant difference between their performance in the two-probability and multiple-probability tasks. All behavioral tasks were performed in a dark sound-attenuated chamber, with white noise delivered between 2–60 kHz.

All surgical and experimental procedures were in accordance with the National Institutes of Health Guide for the Care and Use of Laboratory Animals and approved by the Johns Hopkins University Animal Care and Use Committee. This experimental setup and some analyses of the data for electrophysiological recordings and matching behavior have been reported previously (Bari et al., 2019; Trepka et al., 2021; Bari & Gershman, 2022). All analyses reported in this study are novel.

#### Monkey experiment

In the monkey experiment (Tsutsui et al., 2016), two adult male macaque monkeys (*Macaca mulatta*) weighing 5.5–6.5 kg were trained in a reward-based foraging task. To initiate a trial, the monkeys fixated on the red fixation spot in the center of a computer screen and contacted a touch-sensitive, immobile resting key at elbow height. After fixation, two stimuli (two different visual fractal objects, A and B) appeared simultaneously on the screen to the left and right of the fixation point (5° visual angle). Two stimuli were assigned pseudorandomly to the left and right positions and were not related to reward. To make a choice, each monkey made a saccadic eye movement to the target of its choice within a time window of 0.25–0.75s (**Figure 1B**). Reward was a drop of juice (0.7 ml) delivered via a spout in front of the animal’s mouth.

The reward probabilities of objects A and B were independently calculated in every trial, such that the instantaneous reward probability increased for the unchosen option as a function of the number of consecutive unchosen trials (Eq. 1):

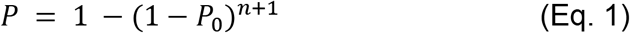

where *P* is the instantaneous reward probability of an object, *P_0_* is an experimentally imposed, baseline reward probability, and *n* is the number of trials that the object had been consecutively unchosen. This equation implies that an unchosen option became more rewarding the longer it remained unselected. The reward probability fell back to the baseline probability with every choice of that object, regardless of whether that choice was rewarded or not. Functionally, this algorithm implements the identical baiting mechanism used in the mouse experiment (also see Eq. 2 in Bari & Gershman, 2022). The baseline reward probabilities (*P_0_*) were chosen from 20 different sets of reward schedules (**Figure S1D**). The two baseline probabilities were held constant for each option for a fixed number of trials (block) and changed across blocks (block switches) such that the previously better option (with higher *P_0_*) became the worse option (lower *P_0_*). The lengths and reward schedules of each block changed randomly from one block to the next, typically ranging from 50 to 150 trials without any cues to the animals. In total, the two monkeys performed 251 blocks during 103 session days (mean 2.43 block switches per session).

All animal procedures conformed to the US National Institutes of Health Guidelines and were approved by the Home Office of the United Kingdom. This experimental setup and some analyses of the electrophysiological recording data have also been reported previously (Tsutsui et al., 2016). All analyses reported in this study are novel.

### Entropy-based metrics

Here, we utilized previously introduced entropy-based metrics to quantify learning and choice behavior (Trepka et al., 2021). This includes entropy of choice strategy (H(str)), conditional entropy of reward-dependent strategy (ERDS), conditional entropy of option-dependent strategy (EODS), mutual information between reward outcome and choice strategy (MIRS), and mutual information between choice and choice strategy (MIOS) (**Figure 1**). Briefly, H(str) measures consistency or randomness in the adopted choice strategy in terms of *stay* or *switch*, whereas ERDS and EODS measure the same consistency but considering reward feedback and/or choice in the previous trial. MIRS, on the other hand, measures the amount of information shared between the adopted strategy (stay or switch) and reward feedback (win or lose). The maximum value for these quantities is equal to 1 corresponding to completely random behavior (equally adopting stay or switch strategy) or no relationship between reward outcome and choice strategy. Conceptually, metrics based on conditional entropy and mutual information aim to capture the degree to which the animal’s choice behavior can be explained or predicted by the information prior to that decision, such as by presence of reward (ERDS, MIRS) or previous selection (EODS, MIOS).

More specifically, ERDS is calculated as follows:

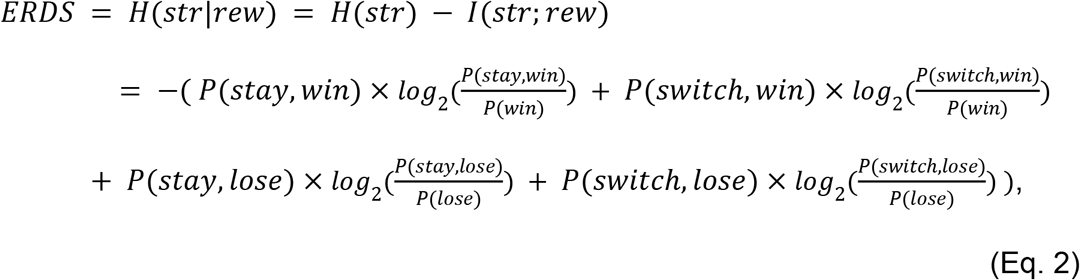

where *str* is strategy coded as stay (1) or switch (0), and *rew* is the previous reward outcome coded as reward (1) or no reward (0). As seen above, the more traditional measures win-stay and lose-switch appear in this equation as and 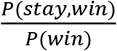 and 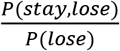, respectively. Additionally, H(str) above is the entropy of strategy (stay or switch), and I(str; rew) is the mutual information of strategy and reward (MIRS):

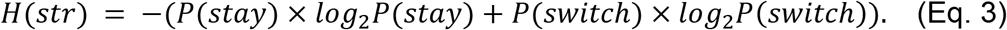

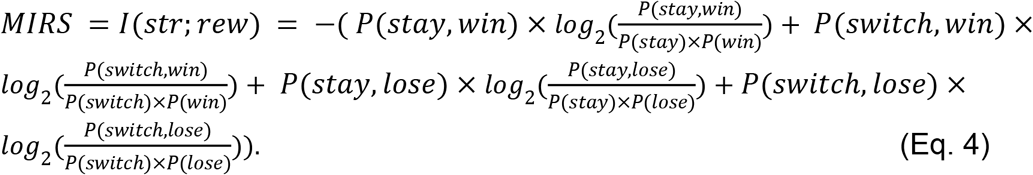

As these terms suggest, ERDS measures the dependence of adopting a response strategy on reward feedback (*win* or *lose*). Lower ERDS values correspond to decreased randomness in the variable and thus more consistency in the utilized strategy.

Similarly, we also used the entropy of option-dependent strategies (EODS), equal to a conditional entropy of stay/switch strategy (*str*) given the choice of better or worse option from the previous trial (*opt*):

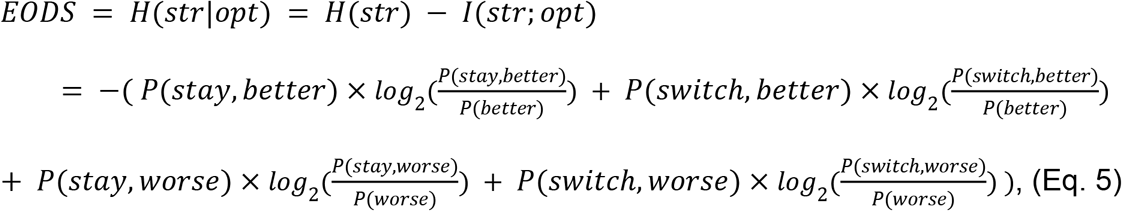

where *opt* is the previous choice in terms of the better or worse option (1 = better option and 0 = worse option), and *I*(*str; opt*) is the mutual information of strategy and option (MIOS):

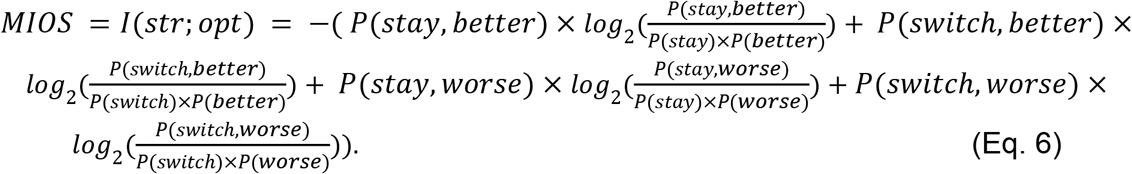

Note that from the above equations, ERDS = H(str) – MIRS and EODS = H(str) – MIOS. This implies that, for example, increased ERDS in one setting could be due to increased H(str) or decreased MIRS, or both.

### Quantifying expected uncertainty and expectation of block switches in the reward environment

In both mouse and monkey experiments, baseline reward probabilities associated with two choice options were fixed within a block of trials, but block lengths changed randomly from one block to the next. This means that within a given block, animals could develop an estimation for uncertainty in reward feedback, often referred to as expected uncertainty corresponding to uncertainty attributable to the stochastic or probabilistic nature of reward outcome (Soltani & Izquierdo, 2019). Additionally, the animals could form an expectation about block lengths or how long reward probabilities remain unchanged. Such expectation could have significant effects on behavior, because the animals could erroneously perceive that a block switch happens after a certain number of trials have passed. Both types of expectation and their effects on behavior might be read from the animal’s responses, such as from the changes in the adopted strategies captured by ERDS or EODS. Therefore, we defined different measures to quantify the different types of expectation as explained below.

#### Expected uncertainty

As the animals performed a wide range of different reward schedules (**Figure S1**), we defined expected uncertainty as the difference between baseline reward probabilities for better (*P_B_*) and worse option (*P_W_*) for each block: Δ*P* = *P_B_* - *P_W_*. For the monkey experiment, base reward probabilities (*P_0_*) for better and worse options were used to compute this metric. The larger the quantity Δ*P*, the easier it is for the animal to discern which is the better option, and hence, experienced less uncertainty in determining the better option. This uncertainty was deemed “expected” in the sense that *P_B_* and *P_W_* were stable for the entire block and reflected stochasticity in the environment. We note quantifying expected uncertainty with Δ*P* is different from the expected uncertainty defined as the overall variance in reward outcome, which can be computed using the average absolute value of reward prediction error (Soltani & Izquierdo, 2019).

Using Δ*P*, we further categorized the blocks into *more uncertain* and *less uncertain* blocks based on the median split of Δ*P* within each subject. Blocks that had larger Δ*P* than the median were categorized as less uncertain, and smaller Δ*P* as more uncertain blocks. We hypothesized that increased uncertainty should result in more stochastic behavior, and therefore *more uncertain* blocks with Δ*P* smaller than median would be characterized by higher entropies or lower mutual information. Overall, the mouse experiment had a median Δ*P* of 0.35 (*M* = 0.330, *SD* = 0.042), and the monkey experiment had median Δ*P* of 0.25 (*M* = 0.281, *SD* = 0.122).

#### Expected block length

As the animals experienced a range of different block lengths, we hypothesized that the animals would keep an expectation of how long the current block is, and that it is updated at every block switch. More specifically, considering *L_i_* (length of block *i*), we calculated the expected block length *E*[*L_i_*] as follows:

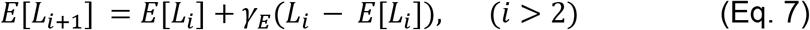

where *γ_E_* is an update rate (for computing block-length expectation) controlling how much to weigh the recent evidence, analogous to the learning rate in the reinforcement learning models. The amount of update depends on the *block length prediction error (BLPE*) defined as the difference between actual and expected block length at block *i*:

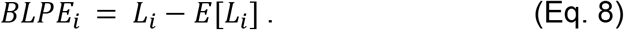

Negative BLPE indicates that a given block was shorter than expected, and positive BLPE suggests that a block was longer than the expected length.

Equation 7 implies that, the larger the *γ_E_* is, the animal weighs the recent evidence more so to update the estimated block length for the next block. For our analysis, we set *γ_E_* = 0.1. For the very first block, *E*[*L_1_*] was set to the median block length of each dataset: *E*[*L_1_*] = 54 and *E*[*L_1_*] = 51 for mice and monkeys, respectively. Any sessions that did not have any block switches were omitted from this update, and *E*[*L_i_*] was held constant for that block, given that it did not contribute to forming an expectation of block switches. Mouse data did not have such single-block sessions, whereas monkey data had 40 blocks that terminated without block switches (15.9% of total blocks).

Alternatively, the expected block length can be defined from a window of size *N*, as the average block length for the past *N* blocks:

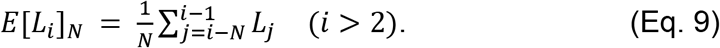

Here, the larger window size *N* corresponds to smaller update rate *γ_E_*, meaning that the animals estimate expected block length from a longer timescale.

Finally, we used *BLPE* (Eq. 8) to categorize blocks as *shorter than expected* (*SE*) or *longer than expected* (*LE*) blocks, and compared behavior across these two types of blocks. We hypothesized that animals could develop an expectation of block switches through experience and use this to modulate their behavior. More specifically, we assumed that if the actual block length (*L_i_*) is past the expected length (*E*[*L_i_*]), the animal would be more likely to incorporate reward feedback as if this feedback signals changes in the environment. In terms of entropy measures, this could be reflected in higher entropies of reward-dependent (ERDS) or option-dependent strategy (EODS), and/or lower mutual information between strategy and reward (MIRS) or previous option (MIOS).

### Reinforcement learning (RL) models and fitting choice data

#### RL models

We fit the choice behavior of mice and monkeys using four standard reinforcement learning models. The first model, which we refer to as RL1, has two parameters: a single learning rate ***α*** and an inverse temperature ***β*** representing choice consistency. The value of the chosen option (*Q_C_*) is updated as follows:

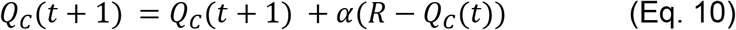

where *C* ∈ {*Left, Right*} for mice and *C* ∈ {*Object A, Object B*} for monkeys, and *R* indicates presence (1) or absence (0) of reward on the given trial *t*.

The second model, referred to as RL1_decay_, has an additional parameter decay rate ***γ***_decay_ for the value of the unchosen option. In this model, the value of the unchosen option (*Q_U_*) decays passively, using the following equation:

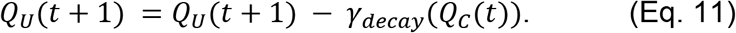

The value estimates for the chosen option were updated as in Eq. 10.

The third and fourth models, RL2 and RL2_decay_, are identical to RL1 and RL1_decay_ except that there are now two separate learning rates for rewarded and unrewarded trials (***α***_+_ and ***α***_-_, respectively). For this set of models, the value of the chosen option (*Q_C_*) was updated as follows:

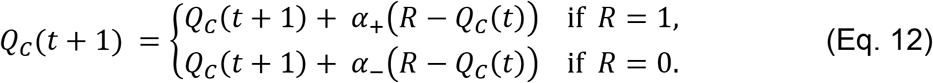

RL2 did not include a decay term, whereas RL2decay followed a decay equation for an unchosen option as in Eq.11.

For all models above, the probability of choosing an outcome was then computed using a softmax function:

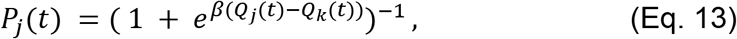

where *j* and *k* correspond to *Left* and *Right* for mice, and *Object A* and *Object B* for monkeys. The minimum and maximum values of ***β*** were set to 1 and 100, respectively.

#### Model fitting process

We used the standard maximum likelihood estimation method to fit and estimate the best-fit parameters for the described models. One set of model parameters was fit to each block of mice and monkeys’ choice data. When fitting and simulating RL models with mouse data, we treated infrequent miss trials (average of 3.64 per session) and no-go trials (average of 16.83 per session) as if they had not occurred. For the very first block of the session only, value estimates for the two options (*Q*) were initialized at 0.5. For the consequent blocks after block switches (within the same session), initial values were set equal to *Q* values from the last trial of the previous block, given by its best-fit parameters. Fitting was performed using the MATLAB optimization function *fmincon*, repeating the search for 10 random parameter values.

To quantify goodness-of-fit, we computed the Akaike Information Criterion (AIC) for each block of the mouse and monkey datasets:

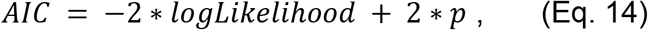

where *p* is the number of free parameters in a given model. Since each block had different number of trials, we further normalized the AIC by the number of trials in each block, i.e., computing AIC per each trial:

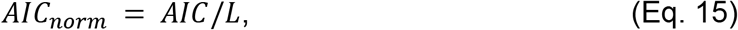

where *L* is the number of trials or length of each block. We determined the winning model as the one with the lowest mean *AIC_norm_* across all blocks.

We further computed the probability that a given model is the best model for the dataset given the set of candidate models (*Akaike weights*). Specifically, Akaike weights (*w_i_*) for the *i*-th model (*M_i_*) in a set of *n* models were computed as follows:

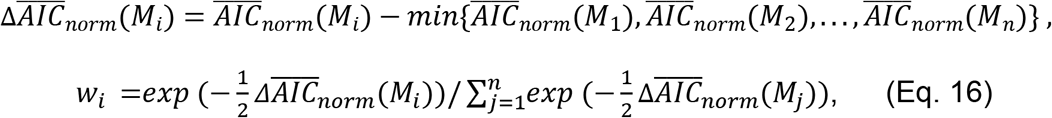

where 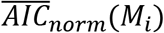 indicates the mean *AIC_norm_* of *M_i_* across all blocks, and 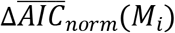 is the difference between the mean *AIC_norm_* of *M_i_* and the minimum mean *AIC_norm_* among all the candidate models.

### Statistical analysis and visualization

Due to log-transformation in the entropy equations, the distributions of most of the entropy-based metrics are highly skewed (**Figure S2**). Accordingly, we primarily employed a nonparametric test: a two-sample Kolmogorov-Smirnov test with a null hypothesis that the two samples are from the same continuous distribution. The test statistic *D* for this test measures the difference between the CDFs of the two distributions.

To avoid confounds with the data size and the computed entropy values, we kept the same bin size (number of trials) from each block for computing the metrics. Specifically, we selected the last *N* trials of each block while ensuring that 10 trials had passed after the start of each block. Blocks that had too few trials to compute reliable entropy values were removed from the analysis. To minimize the number of excluded blocks while keeping its proportion consistent across datasets, we set *N* = 30 for the mouse dataset and *N* = 20 for the monkey dataset. This removed 511 blocks out of 3768 (13.56%) for mice and 36 blocks out of 251 (14.34%) for monkeys.

Finally, as an auxiliary to the Kolmogorov-Smirnov test, we also used a permutation test to compare the entropy-based metrics between two given conditions. More specifically, for each permutation, we randomly assigned blocks (without replacement) into two groups with sample size *n_1_* or *n_2_*, corresponding to the total number of blocks in the two conditions that were being compared. A single entropy-based metric was computed for each type of block by concatenating all trial vectors from the sampled permutation. For consistency, we excluded the same 10 trials after the start of each assigned block. Test statistics for each sample were then computed by taking the difference in metrics between the first and the second condition. We computed the p-value by computing the proportion of sampled test statistics that were as extreme as the observed value.

All data analysis was performed in MATLAB 2020b (Mathworks). All violin plots in the figures were created using an adapted version of a custom toolbox *al_goodplot* in MATLAB (Legouhy, 2022). In the violin plots, the rectangles within the kernel distributions indicate the 25th, 50th (black horizontal line), and 75th percentiles, and circles indicate the mean of the data. For the visualization of the entropy time course, we used confidence intervals from the basic bootstrapping method which computes the difference between observed and sampled test statistics (David & Hinkley, 1997). As mentioned above, to determine statistical significance, a non-parametric test (Kolmogorov-Smirnov test) was primarily used. A parametric t-test was used where appropriate, after verifying that the sample distribution was symmetric and approximately normal. All tests were two sided and alpha was set at 0.05, except otherwise mentioned.

## Results

To study how uncertainty related to different types of regularities in the reward environment affects learning and choice behavior, we used multiple metrics based on information theory (**Figure 1**) to reanalyze data from two dynamic foraging tasks in mice and monkeys. In both tasks, animals were trained to choose between two alternative choice options (left and right port for mice, and two abstract pictures for monkeys; **Figure 2A–B**) in order to obtain reward. Reward, a drop of water for mice or a drop of juice for monkeys, was delivered probabilistically and independently of the unchosen option (see below for more details). After a variable number of trials (a block) the reward probability changed for the two options without any cue to the animals, often resulting in a switch between the better and worse options (sometimes referred to as a reversal). The length of each block changed randomly from one block to the next.

**Figure 2.**
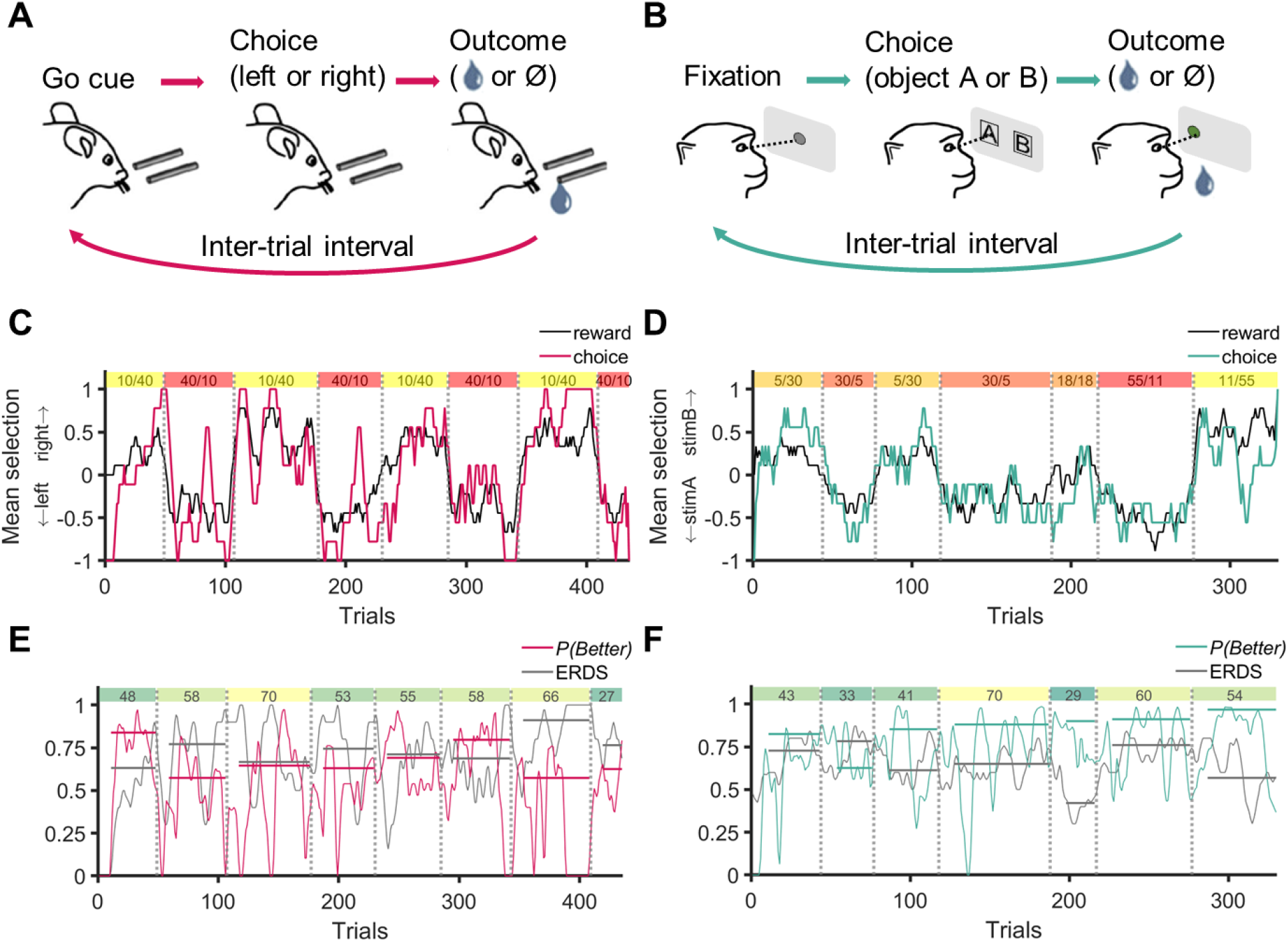
Experimental paradigms and example behavioral results in mice and monkeys. (**A, B**) Timeline of a single trial during experiments in mice (A) and monkeys (B). To initiate a trial, mice received an olfactory go-cue (or no-go cue in 5% of trials), and monkeys fixated on a central point. Animals indicated a choice by licking (mice) or making a saccadic eye movement (monkeys) toward their choice among the two options (left/right port for mice and two abstract pictures for monkeys). Reward (a drop of water for mice and a drop of juice for monkeys) was delivered stochastically based on dynamically assigned reward probability on the chosen option. (**C, D**) Average choice and reward outcomes using a moving window of 10 trials for a representative session in mice (C) and monkeys (D). Numbers in the colored boxes indicate reward probabilities (in %) of two options within each block. (**E, F**) Time course of performance and consistency in reward-dependence strategy in example sessions in mice (E) and monkeys (F). Performance (probability of choosing the better option, P(Better)) and conditional entropy of reward-dependent strategy (ERDS) are computed from a moving window of 10 trials from the same example sessions. Numbers in the colored boxes show the length *L* (number of trials) of each block. Horizontal lines show the overall metrics during each block, computed from all trials in the block (excluding ten trials after block switch).

In the mouse experiment, rewards were baited such that if reward was assigned on a given side and that side was not selected, reward would remain on that side until the next time that side was selected. The monkey experiment had a functionally identical baiting mechanism, where instantaneous reward probability increased for the unchosen option such that selection of the unchosen option became more rewarding as time since that option was chosen increased. The reward probability for a given option fell back to the assigned baseline probability with every choice of that option, irrespective of whether that choice was rewarded or not.

Overall, both mice and monkeys learned reward associations, and their choices closely tracked the reward schedules, reflected in the close match between the probability of choosing the better option, P(Better), and the relative probability of reward for that option (**Figure 2C–D**). As expected, when animals chose the better option more consistently, conditional entropy of reward-dependent strategy (ERDS) was lower (**Figure 2E–F**). This was also reflected as a negative correlation between P(Better) and ERDS (Spearman’s correlation, *r* = −0.647, *p* = 0), option-dependent strategy (EODS; *r* = −0.836, *p* = 0), and baseline entropy of strategy (H(str); *r* = −0.675, *p* = 0). In monkeys, however, the relationship between performance and ERDS was not significantly correlated (*r* = −0.0185, *p* = .788), whereas higher performance was associated with lower EODS only (*r* = −0.400, *p* = 1.20×10^-9^). Interestingly, H(str) was positively correlated with performance (*r* = 0.315, *p* = 2.50×10^-6^) indicating that an overall random strategy facilitated the performance in monkeys.

### Entropy-based metrics capture changes in behavior due to block switches

To examine animals’ response to changes in reward schedule caused by block switches, we next computed different behavioral metrics before and after those events. In order to determine whether the metrics changed significantly after block switches, we used a paired-sample t-test for each pair of metrics around block switches. As expected, the overall performance decreased after block switches for both mice (**Figure 3A**; *t*(2992) = 27.6, *p* = 1.41×10^-149^; Cohen’s *d* = 0.52) and monkeys (**Figure 3D**; *t*(139) = 6.13, *p* = 8.48×10^-9^; Cohen’s *d* = 0.499). The decrease was more pronounced for mice, because before block switches mice exhibited stronger selection of the better side (*M* = 0.744, *SD* = 0.204) than the monkeys’ selection of the better stimulus (*M* = 0.617, *SD* = 0.11).

**Figure 3.**
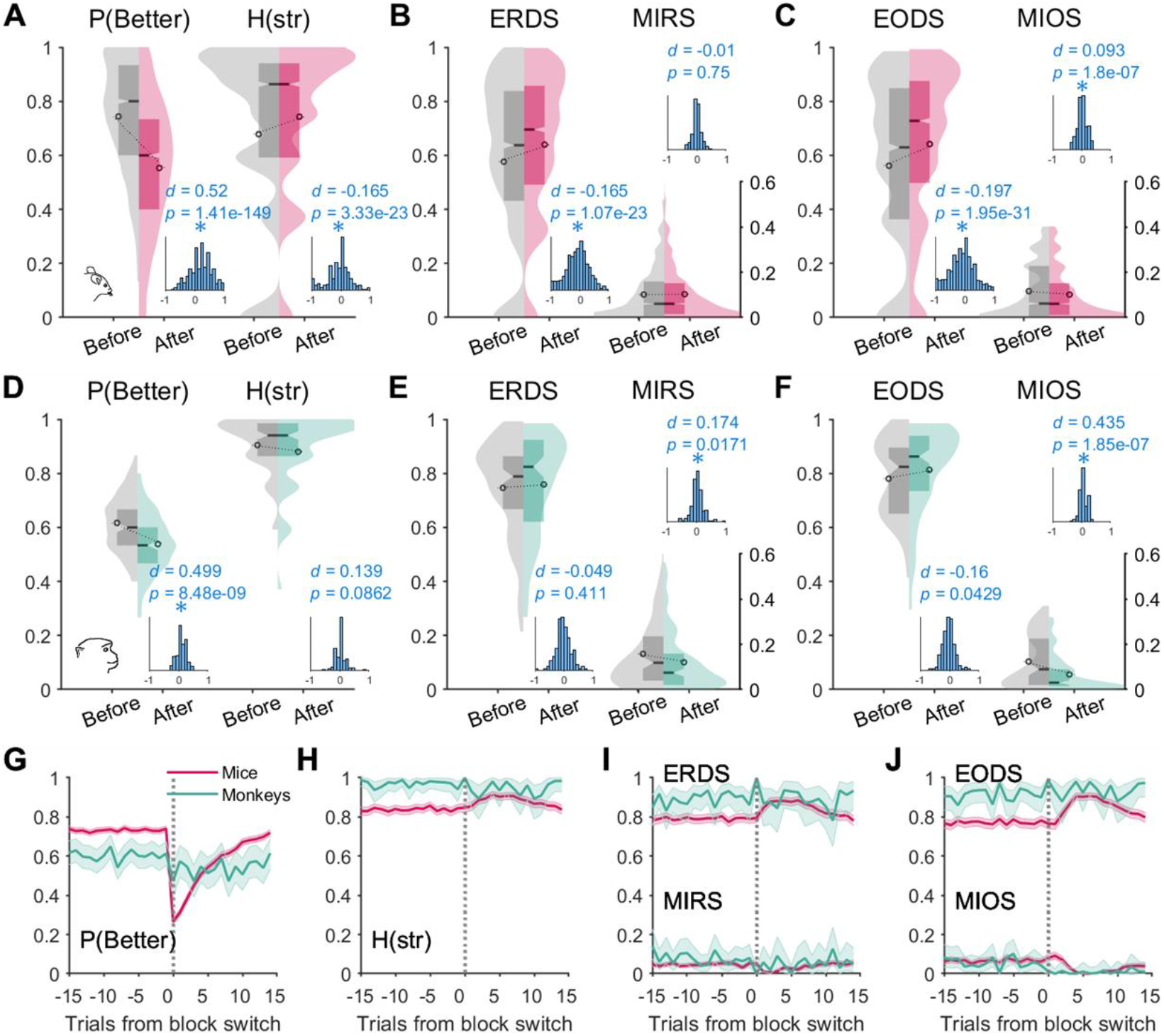
Changes in performance and behavioral metrics around block switches. (**A–F**) Violin plots show distributions of performance and entropy-based metrics computed from 15 trials before and after block switches in mice (A–C) and monkeys (D– F). In the violin plots, the rectangles within the kernel distributions indicate 25th, 50th (median, black horizontal line), and 75th percentiles. Circles indicate the mean of the data. Insets show histograms of paired difference between metrics before and after each block switch. Reported values are Cohen’s *d* and p-values based on paired-sample t-test comparing before and after block switches (two-sided test, corrected for two multiple comparisons). (**G–J**) Time course of different metrics around block switches. Plots show running averages of metrics computed from the same trials in time relative to block switches (vertical dotted line) across all blocks of the experiment. Shaded error bars indicate 95% confidence intervals from bootstrapping method using *N* = 10,000 samples for each trial point. Overall, mice had significantly lower H(str), ERDS, EODS, and MIRS than monkeys.

Comparisons of entropy-based metrics in mice revealed that block switches increased inconsistency in reward-dependent choice strategy (ERDS) (**Figure 3B**; *t*(2992) = −10.12, *p* = 1.07×10^-23^; Cohen’s *d* = −0.165) mainly by increasing the overall entropy in choice strategy (stay vs. switch) measured by H(str) (**Figure 3A**; *t*(2992) = −10.01, *p* = 3.33×10^-23^; Cohen’s *d* = −0.165), as there was no evidence for a change in the link between reward outcome and adopted choice strategy measured by mutual information of reward-dependent choice strategy (MIRS) (**Figure 3B**; *t*(2992) = −0.32, *p* = 0.750; Cohen’s *d* = −0.01). Similarly, mice became less consistent in how they stayed on or switched from the better option measured by EODS (**Figure 3C**; *t*(2992) = −11.8, *p* = 1.95×10^-31^; Cohen’s *d* = −0.197), whereas the link between selected option and adopted choice strategy (measured by MIOS) became weaker after block switches (**Figure 3C**; *t*(2992) = 5.23, *p* = 1.80×10^-7^; Cohen’s *d* = 0.093).

In contrast, comparisons of entropy-based metrics in monkeys showed that block switches only made the link between reward outcome and choice strategy (measured by MIRS) weaker (**Figure 3E**; *t*(139) = 2.41, *p* = .0171; Cohen’s *d* = 0.174) without any changes in reward-dependent choice strategy (**Figure 3E**; *t*(139) = −0.82, *p* = .411; Cohen’s *d* = −0.049) or the overall choice strategy, H(str) (**Figure 3D**; *t*(139) = 1.73, *p* = .0862; Cohen’s *d* = 0.139). Similarly, the link between the selected option and adopted choice strategy (measured by MIOS) became weaker right after block switches (**Figure 3F**; *t*(139) = 5.49, *p* = 1.85×10^-7^; Cohen’s *d* = 0.435) with a noticeable but not statistically significant change in option-dependent choice strategy, EODS (**Figure 3F**; *t*(139) = −2.04, *p* = .0429; Cohen’s *d* = −0.16).

The similar and contrasting effects of block switches in the two species could also be seen from the time course of behavioral metrics over time (**Figure 3G–J**). Comparing the consistency of choice behavior in mice and monkeys around block switches, we observed that the monkeys’ behavior was more random and less predictable by previous reward outcome or choice. Kolmogorov-Smirnov tests between mouse and monkey data revealed significantly higher H(str) (**Figure S2D**; *D* = 0.441, *p* = 5.15×10^-35^), ERDS (**Figure S2B**; *D* = 0.276, *p* = 5.85×10^-1014^), and EODS (**Figure S2C**; *D* = 0.361, *p* = 1.19×10^-23^) in monkeys. Although MIOS did not significantly differ between the two species (**Figure S2F**; *D* = 0.0944, *p* = 0.0517), MIRS was slightly higher in monkeys (**Figure S2E**; *D* = 0.231, *p* = 5.93×10^-10^). These opposing patterns of MIRS and ERDS across species can be explained by noticing that, compared to mice, the monkeys’ larger shared information between reward outcome and subsequent choice strategy (measured by MIRS) was not strong enough to result in more consistent reward-dependent strategy (measured by ERDS), perhaps due to an overall more random choice strategy (H(str)).

Comparing the entropy metrics within each species using paired t-test, we found that mice exhibited an option-dependent strategy more so than a reward-dependent one: EODS tended to be lower than ERDS (*t*(3226) = 11.49, *p* = 5.48×10^-30^; Cohen’s *d* = 0.202), and MIOS tended to be higher than MIRS (*t*(3226) = −11.49, *p* = 5.48×10^-30^; Cohen’s *d* = −0.202). In contrast, monkeys showed more of a reward-dependent strategy compared to an option-dependent one, as indicated by lower ERDS than EODS (*t*(214) = −2.36, *p* = 0.0193; Cohen’s *d* = −0.161) and higher MIRS than MIOS (*t*(214) = 2.36, *p* = 0.0193; Cohen’s *d* = 0.161).

This pattern of the adopted strategy was consistent across individuals within each species. For mice, thirteen individuals (72.2% of total) had a lower average EODS than ERDS, and for eight of these individuals this effect was statistically significant (two-sided paired t-test, p < .05; **Figure S3A**). For monkeys, both individuals had a lower average ERDS than EODS, but only one showed a statistically significant difference (p = .0206 and p = .4980; **Figure S3B**). We further addressed whether the adopted strategy was also consistent across sessions for each individual by examining how the difference between ERDS and EODS changed as a function of time. Negative values of (ERDS - EODS) suggest preference for a reward-dependent strategy, whereas positive values would suggest preference for an option-dependent strategy. For each individual, we regressed these quantities by the number of sessions completed (**Figure S3C–D**). For mice, we found that twelve individuals (66.7% total) showed positive regression coefficients, with four for which the effect was statistically significant (Figure S3C). In contrast, six individuals (33.3%) showed a negative fitted slope, with two for which the effect was statistically significant. For monkeys, both individuals showed a negative slope, but none were statistically significant (**Figure S3D**). These results suggest that, through time, the majority of mice preferred to adopt an option-dependent strategy, whereas monkeys tended toward a reward-dependent strategy.

### Effects of environmental regularities and uncertainty on behavior

One of the main aims of this study was to investigate how animals’ choice behavior is affected by regularities and accompanied uncertainty in the reward environment. To answer this question, we first ran exploratory regression analyses to predict behavioral metrics based on a few experimental parameters (**Figures S4, S5**). This included the difference in reward probabilities for the better and worse choice options (*P_B_* and *P_W_*) reflecting uncertainty in determining the better option, the sum of reward probabilities measuring richness of the reward environment, and the block length in the previous and current blocks, aiming to measure whether animals developed an expectation about block lengths and the stability of reward environment. We note, however, that regression may not be the most suitable analysis because of the non-linear nature of entropy-based metrics (**Figure S2**). Nevertheless, these regression analyses pointed to the effects of a difference in reward probabilities and block length on multiple entropy-based metrics, indicating that certain regularities in the environment affected learning and choice behavior.

To better understand these effects, we next grouped blocks based on the block length and the difference in reward probabilities and computed different metrics for each group of blocks. First, we divided blocks based on the difference in reward probabilities for the better and worse options and categorized blocks into *more uncertain* and *less uncertain* blocks based on the median split of *P_B_* - *P_W_* in each subject (see **Methods** for more details). The difference in reward probabilities measures uncertainty in determining the better choice option. Blocks that had larger *P_B_* - *P_W_* than the median were categorized as *less uncertain* and those with smaller *P_B_* - *P_W_* as *more uncertain* blocks. Second, we also divided blocks depending on whether the current block length was longer than expected or not. More specifically, we assumed that animals developed an expectation of block switches by estimating a value for the length L of a block at each block *i*, denoted *E*[*L_i_*], updated at every block switch. This update was done using the *block length prediction error (BLPE*) equal to the difference between the actual and expected block lengths (see **Methods** for more details). Negative BLPE corresponds to a block that was *shorter than expected* (*SE* blocks) based on the animal’s experience, whereas positive BLPE indicates that a block was *longer than expected* (*LE* blocks).

With regard to uncertainty in determining the better option, our hypothesis was that increased uncertainty should result in slower learning and/or more stochastic behavior. In addition, we hypothesized that when blocks are longer than expected (LE blocks), choice or adopted choice strategy would be more random as animals expect changes in reward probabilities more strongly and in the absence of any changes in the environment. Finally, expectation of block switches could interact or depend on the level of uncertainty in the environment.

Supporting these hypotheses, we found that entropy-based metrics were systematically affected by both uncertainty and expectation of block switches, but differently for mice and monkeys. In mice, entropy of choice (H(str); **Figure 4A**), reward-dependent choice strategies (ERDS; **Figure 4B**), and option-dependent choice strategies (EODS; **Figure 4C**) were significantly larger in the more uncertain environments, with no significant difference between mutual information (MIRS or MIOS; **Figure 4B, C**). Considering the link between entropies, conditional entropies and mutual information (ERDS = H(str) – MIRS and EODS = H(str) – MIOS), these results suggest that the observed increase in H(str) could be due to changes in the effects of reward feedback on choice. In monkeys, however, there was a trend toward a decrease (instead of increase) in the entropy of choice (H(str)), suggesting that the animals’ strategy became more consistent when reward probabilities were closer to each other; i.e., when the environment was more uncertain (**Figure 5A;** Kolmogorov-Smirnov test; *D* = 0.277, *p* = .0250). Post-hoc analysis revealed that this decrease in H(str) in monkeys was due to a higher probability of switching in the more uncertain environment (*M* = 0.668, *SD* = 0.11) compared to the less uncertain one (*M* = 0.508, *SD* = 0.142). Therefore, monkeys increased their exploration in more uncertain environments.

**Figure 4.**
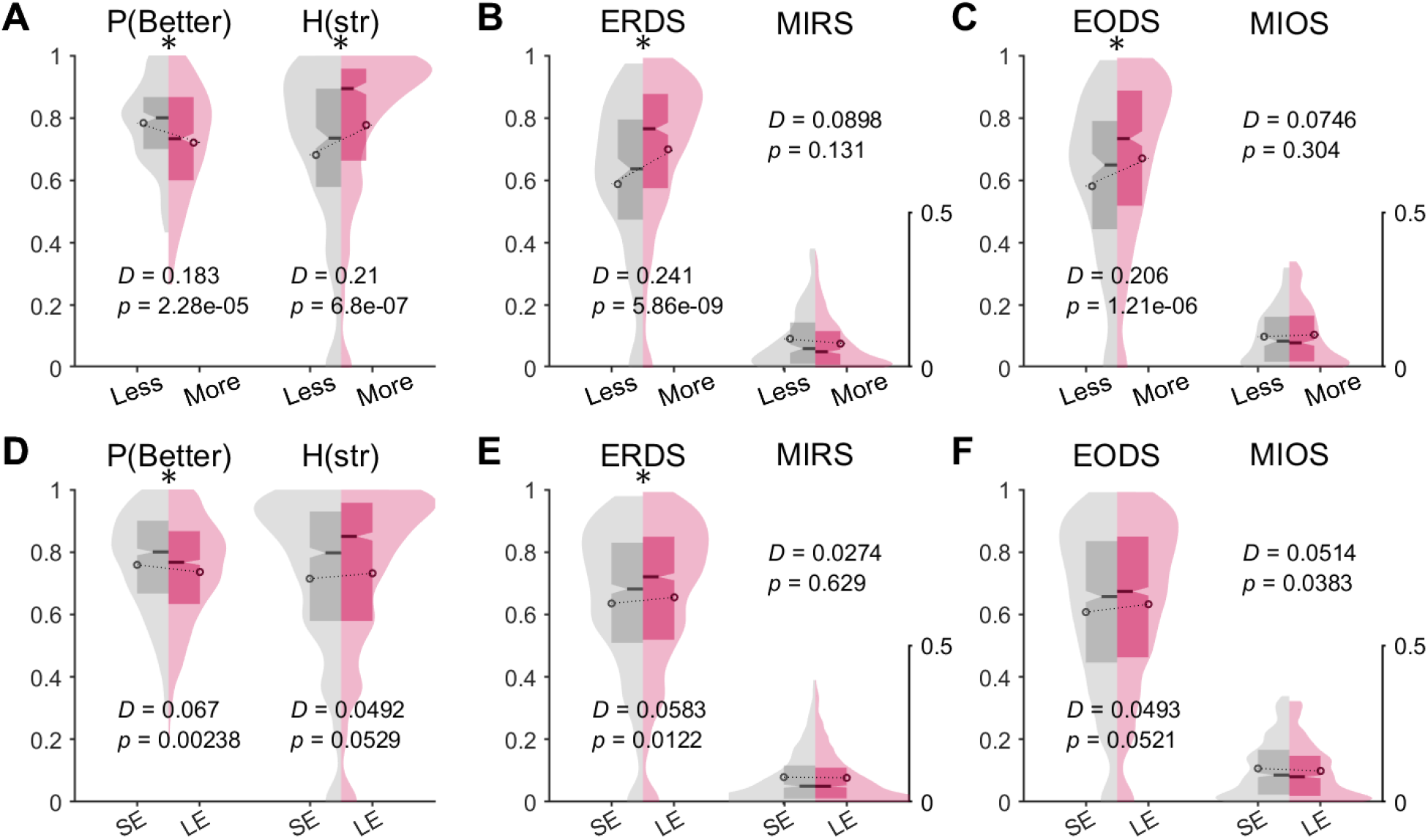
Effects of uncertainty and expectation of block switches on choice behavior in mice. (**A–C**) Effects of uncertainty on choice behavior. Violin plots show the distributions of performance and entropy-based metrics computed from different types of trials according to uncertainty of the reward schedule. All metrics are computed using the last 30 trials in each block. Asterisks show results of two-sample Kolmogorov-Smirnov test with a null hypothesis that the two samples are from the same continuous distribution (two-sided test, corrected for two multiple comparisons). More uncertainty results in more inconsistent choice driven by more stochasticity. (**D–F**) Effects of block switch expectation on choice behavior. SE and LE indicate shorter than and longer than expected blocks, respectively, corresponding to trials before and after the animal expected a block switch. Violin plots show the distributions of performance and entropy-based metrics computed from different types of trials according to block length prediction error. Overall, mice adopted less consistent choice strategies when blocks were longer than expected, especially in more uncertain environments.

**Figure 5.**
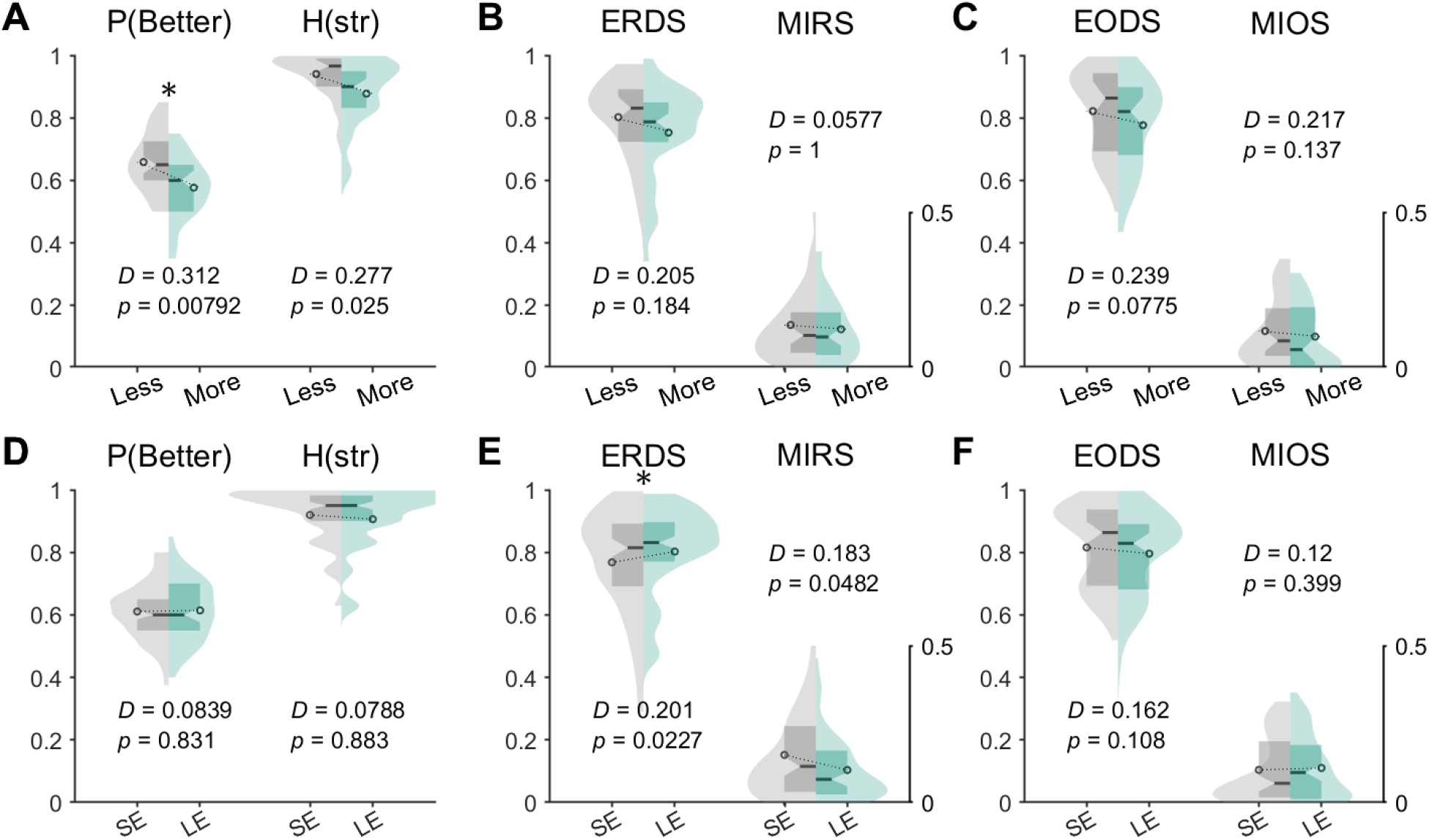
Effects of uncertainty and expectation of block switches on choice behavior in monkeys. (**A–C**) Effects of uncertainty on choice behavior. Violin plots show the distributions of performance and entropy-based metrics computed from different types of trials according to uncertainty of the reward schedule. All metrics are computed using the last 20 trials in each block. Asterisks show results of two-sample Kolmogorov-Smirnov tests with a null hypothesis that the two samples are from the same continuous distribution. (**D–F**) Effects of block switch expectation on choice behavior. SE and LE indicate shorter than and longer than expected blocks, respectively, corresponding to trials before and after the animal expected a block switch. Violin plots show the distributions of performance and entropy-based metrics computed from different types of trials according to block length prediction error. Overall, monkeys adopted less consistent reward-dependent choice strategies when blocks were longer than expected, especially in more uncertain environments.

With regard to expectation of block switches, we found that in mice, ERDS was significantly increased for LE compared to SE blocks (**Figure 4E**; Kolmogorov-Smirnov test; *D* = 0.0583, *p* = .0122). H(str) and EODS followed a similar trend, but the increase was not significant (Kolmogorov-Smirnov test; H(str): *D* = 0.0492, *p* = .0529; EODS: *D* = 0.0493, *p* = .0521; **Figure 4D, F**). The changes in mutual information (MIRS, MIOS) were small or insignificant (**Figure 4E, F**). The effect of expectation of block switches on ERDS in the absence of any clear changes in H(str) and MIRS suggests multiple forms of adjustments including changes in learning and stochasticity in choice. Similarly, expectation of block switches increased ERDS in monkeys (Kolmogorov-Smirnov test; *D* = 0.201, *p* = .0227; **Figure 5D**), and this was accompanied with a moderate but not significant decrease in MIRS (Kolmogorov-Smirnov test; *D* = 0.183, *p* = .0482; **Figure 5D**). Another post-hoc analysis looking at win-stay/lose-switch tendencies of monkeys revealed that the observed change in reward-dependent strategy mainly resulted from reduced probability of win-stay from SE (*M* = 0.627, *SD* = 0.22) to LE blocks (*M* = 0.564, *SD* = 0.196). Lose-switch tendency was overall higher than win-stay but was largely the same between SE (*M* = 0.738, *SD* = 0.158) and LE blocks (*M* = 0.733, *SD* = 0.156). In other words, monkeys adjusted their reward-dependent strategy to expected block switches by switching more from the previously rewarded option.

Finally, to test possible interactions between uncertainty and reward expectation, we further divided SE and LE blocks according to uncertainty (more or less uncertain) and found that the major effect of block switch expectation was mediated by uncertainty in the reward probability. We found that the differences between SE and LE blocks were mainly significant in more uncertain blocks but not in less uncertain ones (**Figure S6**). This suggests that expectation was more pronounced in more uncertain reward environments because in less uncertain environments, reward feedback could signal changes in the reward schedule more easily.

To compare behavioral changes to actual and expected block switches, we performed further analyses focusing on the LE blocks, which allowed us to examine changes to expected block switches. These blocks were longer than expected (L > E[L]) and therefore had expected block switches situated *prior* to the actual block switches. Accordingly, we computed the changes in the behavioral metrics after expected block switches using paired-sample t-tests, similar to the analyses presented in **Figure 3**.

We found that mice reacted to actual and expected block switches similarly; both of these switches were accompanied with increased H(str), ERDS, and EODS, and decreased MIOS (**Figure S7**). Among these metrics, EODS showed the largest changes as suggested by the effect sizes (**Figure S7C**; actual block switch: Cohen’s *d* = −0.209, *p* = 6.37×10^-14^; expected block switch: Cohen’s *d* = −0.225, *p* = 7.24×10^-24^). Such significant effects on both EODS and MIOS suggest that the observed adjustment in behavior after perceived block switches could be mainly due to changes in the option-dependent strategy, as the animals anticipate a switch between the better and worse choice options. Monkeys also similarly reacted to actual and expected block switches with decreased MIRS (**Figure S8E**; actual block switch: Cohen’s *d* = 0.322, *p* = .0131; expected block switch: Cohen’s *d* = 0.346, *p* = .00896). ERDS increased after expected block switches (**Figure S8B**, right; Cohen’s *d* = −0.411, *p* = .00275) more so than after actual block switches (**Figure S8B**, left; Cohen’s *d* = −0.183, *p* = .197).

These results indicate that monkeys adapted to perceived block switches mainly through changes in a reward-dependent strategy, contrary to mice which mostly showed changes in an option-dependent strategy. Overall, the means of paired differences did not significantly differ between actual and expected switches for most of the metrics, as determined by independent-sample t-test even when using the most liberal alpha level at 0.05 (insets in **Figure S7, S8**). Therefore, the above analyses based on behavioral metrics revealed that both mice and monkeys showed consistent adjustments to expected and actual block switches, but monkeys also exhibited some discrepancies in their adjustments (e.g., MIOS in **Figure S8F** inset).

An additional analysis suggests that some of the observed differences in the adopted strategy by mice and monkeys (option-dependent vs. reward-dependent) can be attributed to differences in tasks. More specifically, we examined whether the difference between ERDS and EODS (ERDS - EODS) in a given block of trials depends on the level of uncertainty in determining the better option using Δ*P* (similar to the top rows of **Figures 4** and **5**). We found that for mice, (ERDS - EODS) significantly decreased from less uncertain to more uncertain environments (*M*±*SD*: 0.031 ±0.108 vs. 0.016 ±0.13; two-sided t-test, *p* = .000266; Cohen’s *d* = 0.129; **Figure S9A**). Similarly, monkeys showed a decrease in (ERDS - EODS) between less and more uncertain environments but this decrease was not statistically significant (*M±SD:* −0.022 ±0.144 vs. −0.026 ± 0.139; two-sided t-test, *p* = .862; Cohen’s *d* = 0.0274; **Figure S9B**). Overall, these results raise the possibility that more difficult reward schedules influence animals to tend toward a more reward-based strategy.

Computation of expected block length (*E*(L)) and resulting block length prediction error (BLPE) depend on the parameter *γ_E_* that controls update of *E*(L) (see Eq. 7 in **Methods**). In all of the above analyses related to the effect of block switch expectation, we set *γ_E_* = 0.1. Nonetheless, we also tested whether and how observed differences in entropy-based metrics between SE and LE blocks depend on the value of *γ_E_* (**Figures S10** and **S11**). Although we found differences for a wide range of *γ_E_* values, they tended to be more pronounced for smaller values of *γ_E_*, corresponding to integration of block length over longer timescales. This observation is in line with the intuition that capturing regularities of the environment requires estimation based on multiple block switches.

Finally, we conducted an additional permutation test to compare entropy-based metrics across conditions (*less* vs. *more uncertain*, or *SE* vs. *LE*; see **Methods** for more details). This was done to avoid a possible confound due to small sample size for computation of entropy-based metrics that causes bias toward smaller values (though this bias should impact all conditions similarly as equal numbers of trials were used for different conditions). Overall, results from the permutation test were consistent with and in some cases more likely to be significant than those based on the Kolmogorov-Smirnov test. As for the effect of reward uncertainty in mice (**Figure 4A–C**), we confirmed the same increase in H(str) (**Figure S12A**), ERDS (**Figure S12C**), and EODS (**Figure S12G**), with additional significant decrease in MIRS in more uncertain blocks (**Figure S12E**). The added significant effect of MIRS supports our above interpretation that the observed increase in H(str) is most likely due to the changes in the effects of reward feedback on choice strategy. For the effect of uncertainty in monkeys, we found that H(str) (**Figure S12B**), ERDS (**Figure S12D**), EODS (**Figure S12H**), and MIOS (**Figure S12J**) were all significantly smaller in more uncertain environments. The results of the Kolmogorov-Smirnov test were in the same direction but not significant (**Figure 5A–C**). With regard to the effect of block switch expectation, mice had significantly larger EODS (**Figure S13G**) and smaller MIOS (**Figure S13I**) in LE blocks, whereas monkeys showed significant reduction in MIRS only (**Figure S13F**). These results are consistent with the above findings that mice react to perceived block switches through changes in option-dependent strategy (**Figure S7**), whereas monkeys react through changes in reward-dependent strategy (**Figure S8**).

### Reinforcement learning models reveal underlying mechanisms in response to different types of uncertainty

To reveal possible mechanisms by which uncertainty and expectation of block switches affect choice behavior and learning, we used simple RL models (RL1, RL1_*decay*_, RL2, RL2_*decay*_) to fit data in different blocks of the experiments as described above (**Methods**). The first two models had a single learning rate (RL1), while the other two had separate learning rates for rewarded and unrewarded trials (RL2). We also tested how the fit of data can be improved by including forgetting in the estimated value for the unchosen option or action (captured by a decay rate ***γ*** in RLdecay models).

We found the winning model for fitting mouse data to be RL1_*decay*_ that has a single learning rate ***α***, an inverse temperature ***β***, and a decay rate ***γ*** (**Table 1**). The normalized AIC by number of trials (AIC_norm_) for this model was significantly lower than that of the second-best model, RL2_*decay*_ (paired-sample t-test, *t*(3767) = 12.67, *p* = 5.21×10^-36^; Cohen’s *d* = 0.2063). For the monkey data, the best model was the simplest model, RL1, however, the AIC_norm_ for this model was not significantly lower than that of the second-best model, RL2 (paired-sample t-test, *t*(250) = 1.41, *p* = 0.161; Cohen’s *d* = 0.089). Overall, we found that a single-learning rate model provided a better fit to both species than separate-learning rate models. Mouse data tended to prefer models with a decay term. In contrast, monkeys’ choice data were better fit with models without decay, suggesting a different form of reward probability estimation from mice.

**Table 1.**
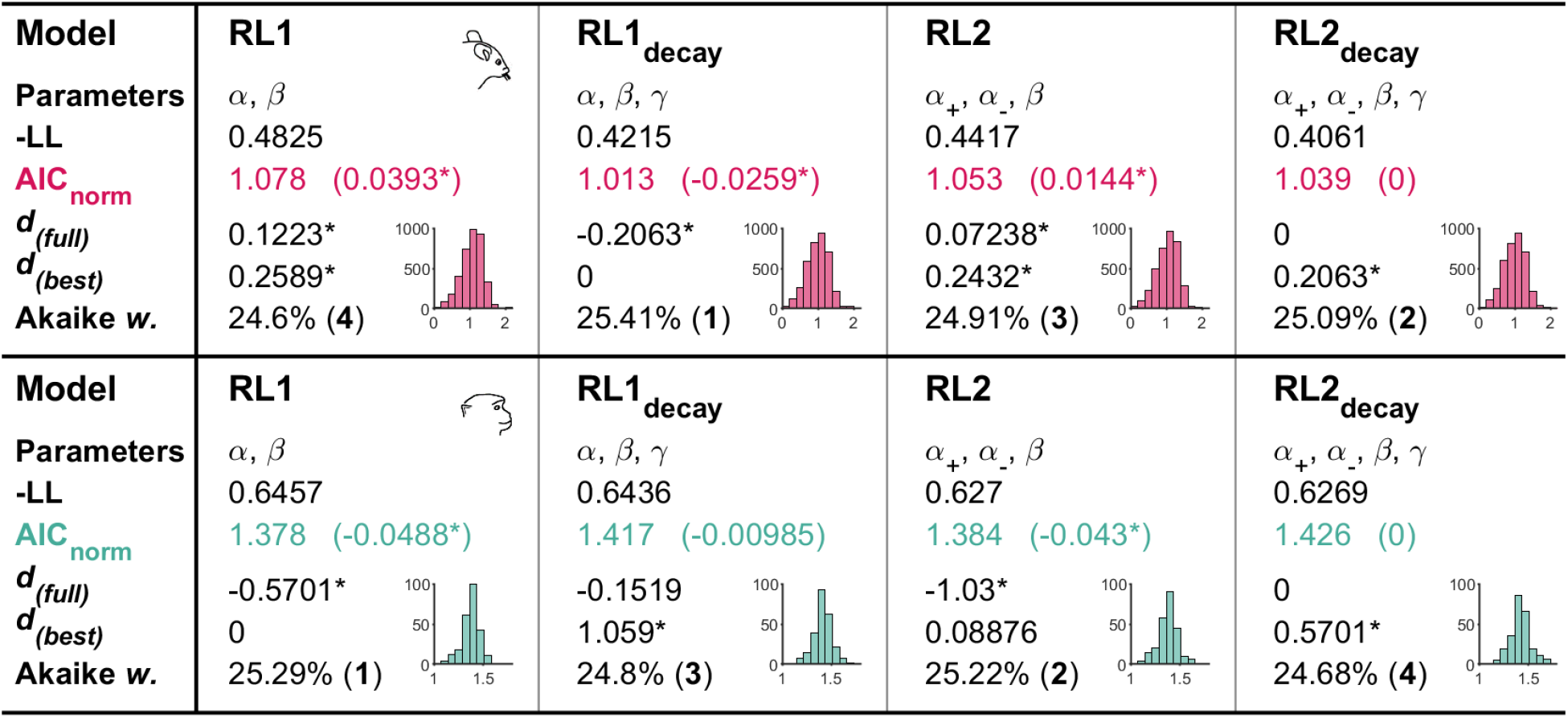
Reinforcement learning models used to fit choice data, their parameters, and their goodness-of-fit. Reported are parameters of a given model and goodness-of-fit, given by negative log-likelihood (-LL) and AIC normalized by the number of trials in each block (AIC_norm_). Values reported in parentheses next to AIC_norm_ are the difference in AIC_norm_ of a given model and the full model (RL2_decay_). Asterisks indicate the significance of this difference indicated by paired-sample t-test (*p* < 0.05/6, multiple comparison corrected). *d*(*full*) indicates Cohen’s *d* value from paired-sample t-tests between a given model and the full model (RL2_decay_), with the asterisk indicating significance. *d*(*best*) indicates Cohen’s *d* value of paired-sample t-tests between a given model and the best model determined by the lowest AIC_norm_. Akaike *w*. = Akaike weights, indicating the probability that a given model is the best model. Integers in parentheses are ranks of a given model. Insets show histograms of AIC_norm_ from the mouse (top row, red) and monkey data (bottom, teal).

| Model | RL1 | RL1<sub>decay</sub> | RL2 | RL2<sub>decay</sub> |
| --- | --- | --- | --- | --- |
| Parameters | $\alpha, \beta$ | $\alpha, \beta, \gamma$ | $\alpha_+, \alpha_-, \beta$ | $\alpha_+, \alpha_-, \beta, \gamma$ |
| -LL | 0.4825 | 0.4215 | 0.4417 | 0.4061 |
| AIC<sub>norm</sub> | 1.078 (0.0393*) | 1.013 (-0.0259*) | 1.053 (0.0144*) | 1.039 (0) |
| $d_{(full)}$ | 0.1223* | -0.2063* | 0.07238* | 0 |
| $d_{(best)}$ | 0.2589* | 0 | 0.2432* | 0.2063* |
| Akaike w. | 24.6% (4) | 25.41% (1) | 24.91% (3) | 25.09% (2) |
| Model | RL1 | RL1<sub>decay</sub> | RL2 | RL2<sub>decay</sub> |
| Parameters | $\alpha, \beta$ | $\alpha, \beta, \gamma$ | $\alpha_+, \alpha_-, \beta$ | $\alpha_+, \alpha_-, \beta, \gamma$ |
| -LL | 0.6457 | 0.6436 | 0.627 | 0.6269 |
| AIC<sub>norm</sub> | 1.378 (-0.0488*) | 1.417 (-0.00985) | 1.384 (-0.043*) | 1.426 (0) |
| $d_{(full)}$ | -0.5701* | -0.1519 | -1.03* | 0 |
| $d_{(best)}$ | 0 | 1.059* | 0.08876 | 0.5701* |
| Akaike w. | 25.29% (1) | 24.8% (3) | 25.22% (2) | 24.68% (4) |

Accordingly, we analyzed the fitted parameters from the winning model of each species (RL2_*decay*_ and RL1 for mice and monkeys, respectively). We hypothesized that a smaller learning rate could be advantageous in more uncertain environments, because it allows animals to obtain a more precise estimate of reward probability in such environments (cf. Figure 1B in Farashahi et al., 2017a). With regard to the expectation of block switches, we hypothesized that when blocks are longer than expected (LE), the increased randomness in choice behavior would result from an increase in the stochasticity of choice and/or decrease in learning, as suggested by results in the previous section. Higher stochasticity in choice can be reflected in the lower inverse temperature ***β***. In contrast, decreased learning could be mediated by decreased learning rate and/or faster decay rate for the unchosen option, both of which could reduce the effects of previous reward feedback on choice behavior.

We found that in mice, the learning rate (***α***) was significantly reduced from less- (*M* = 0.372, *SD* = 0.252) to more-uncertain (*M* = 0.278, *SD* = 0.249) blocks, supporting our hypothesis (**Figure 6A**). We also observed a significantly smaller decay rate (***γ**_decay_*), corresponding to less forgetting of the reward estimate for the unchosen option in more uncertain environments (**Figure 6C**). Overall, these results indicate that more uncertainty resulted in slowing down of value updates in mice. In contrast, the effect of block switch expectation was mediated mostly by changes in the inverse temperature parameter (**Figure 6E**). Specifically, when blocks were longer than expected (LE), the inverse temperature ***β*** was significantly reduced corresponding to more stochasticity in choice (Kolmogorov-Smirnov test, *D* = 0.129, *p* = 4.85×10^-14^). At the same time, longer than expected blocks were accompanied by decreased learning rate and increased decay rate (**Figure 6D–E**). These opposing effects on value updates are interesting as they suggest that when expecting block switches, mice slowed down learning from chosen option while speeding up the forgetting of the unchosen option.

**Figure 6.**
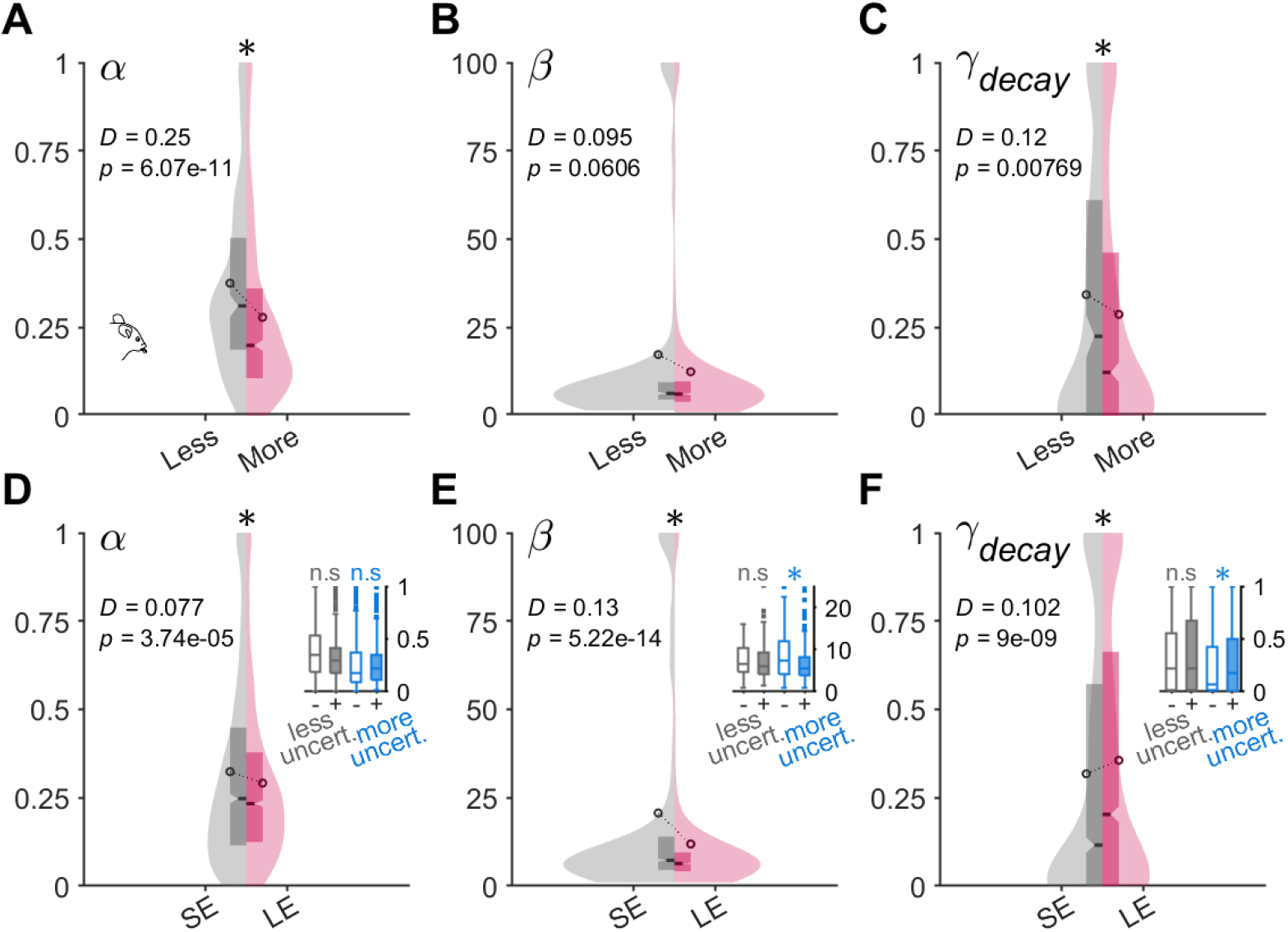
Mechanisms underlying effects of uncertainty and expectation of block switches revealed by fitting of mouse data by reinforcement learning models. Plots show distributions of fitted parameters from RL1_*decay*_ model to mice choice data, categorized by the level of uncertainty (**A–C**) or expectation of block length (**D–F**). ***α*** = learning rate for rewarded and unrewarded trials; ***β*** = inverse temperature or choice consistency; ***γ**_decay_* = decay rate in value estimate for the unchosen option. SE: shorter than expected, LE = longer than expected blocks. Asterisks show results from a two-sample Kolmogorov-Smirnov test. All tests are two-sided, corrected for multiple comparisons of three (number of free parameters). Box plots in the insets show further division of the data into less (gray) or more (blue) uncertain blocks. Asterisks are results of two-sample Kolmogorov-Smirnov tests, corrected for multiple comparisons of six. Interaction effect also exists for the significant RL parameters. Overall, mice responded to the more uncertain reward environment by decreasing the learning rate and slower decay rate for unchosen value. In addition, they responded to trials after an expected block switch by decreasing learning rate, increasing in stochasticity in choice, and having faster decay of unchosen value.

Furthermore, dividing LE and SE blocks into more or less uncertain conditions, we found the same mediating effect of uncertainty as revealed by behavioral metrics. That is, significant changes in the estimated model parameters due to block switch expectation were only observed for more-uncertain blocks, consistent with the interpretation that expectation is more pronounced in more uncertain environments. More specifically, the reduction in ***β*** for LE blocks was significant only for the more uncertain environment (blue bars in **Figure 6E** inset; Kolmogorov-Smirnov test, *D* = 0.219, *p* = 6.17×10^-8^) but not for the less uncertain one (gray bars in **Figure 6E** inset; Kolmogorov-Smirnov test, *D* = 0.105, *p* = .437). Similarly, an increase in ***γ**_decay_* for LE blocks was only observed for more uncertain blocks (blue bars in **Figure 6F** inset; Kolmogorov-Smirnov test; *D* = 0.16, *p* = .00018) but not for less uncertain blocks (gray bars in **Figure 6F** inset; Kolmogorov-Smirnov test, *D* = 0.10, *p* = .503). Interestingly, the learning rate did not show this pattern as there was no significant decrease for LE blocks in less uncertain (gray bars in **Figure 6D** inset; Kolmogorov-Smirnov test, *D* = 0.191, *p* = .0152) or more uncertain environments (blue bars in **Figure 6D** inset; Kolmogorov-Smirnov test, *D* = 0.118, *p* = .0151). This suggests a modulatory effect of local uncertainty about the better option on uncertainty about the expectation of block switches (which could take longer to estimate) via changes in stochasticity in choice and the decay of value estimates for the unchosen option.

In monkeys, we found evidence that greater uncertainty about the better option resulted in significant changes in the estimated model parameters. More specifically, mean learning rate was reduced from less (*M* = 0.443, *SD* = 0.35) to more uncertain (*M* = 0.381, *SD* = 0.33) environments, similar to what we found in mice (**Figure 7A**; Kolmogorov-Smirnov test, *D* = 0.166, *p* = .273). In contrast to mice, however, the effect of block switch expectation was reflected in a significant decrease in the learning rate only (**Figure 7C**; Kolmogorov-Smirnov test, *D* = 0.193, *p* = .0204). This suggests that monkeys slowed down learning, perhaps because around block switches reward feedback was not reliable. Similar to mice, the effects on the learning were more pronounced in less uncertain environments (gray bars in **Figure 7C** inset; Kolmogorov-Smirnov test, *D* = 0.323, *p* = .0657) compared to more uncertain ones (blue bars in **Figure 7C** inset; Kolmogorov-Smirnov test, *D* = 0.275, *p* = .123), although not statistically significant. Inverse temperature was not affected by expectation of block length (**Figure 7D**; Kolmogorov-Smirnov test, *D* = 0.072, *p* = .908). Overall, these results demonstrate that mice and monkeys respond to regularities and uncertainty in the reward environment via different mechanisms.

**Figure 7.**
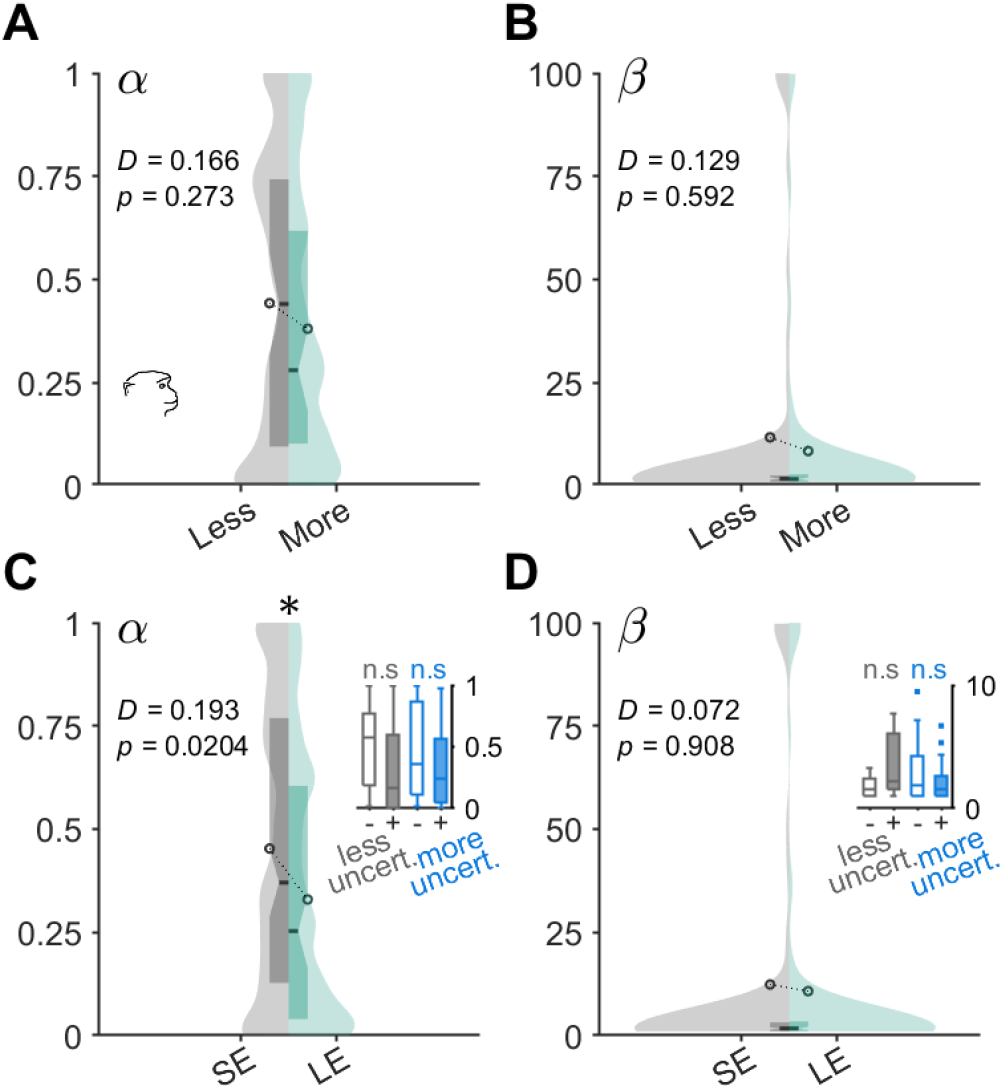
Mechanisms underlying effects of expectation of block switches revealed by fitting of monkey data by reinforcement learning models. Plots show distributions of fitted parameters from RL models to mice choice data, categorized by the effect of expected uncertainty (**A–B**) or expectation of block length (**C–D**). ***α*** = learning rate for rewarded and unrewarded trials; ***β*** = inverse temperature or choice consistency. SE: shorter than expected, LE = longer than expected blocks. Asterisks show results from a two-sample Kolmogorov-Smirnov test. All tests are two-sided, corrected for multiple comparisons of two (number of free parameters). (**A–B**) No parameters were significantly affected by expected uncertainty of the reward environment, similar to the entropy measures. (**C–D**) Monkeys responded to trials after an expected block switch by decreasing learning rate. Box plots in the insets show further division of the data into less (gray) or more (blue) uncertain blocks. Asterisks are results of two-sample Kolmogorov-Smirnov tests, corrected for multiple comparisons of four. Interaction effect was not observed for the significant RL parameters.

## Discussion

How the brain learns from regularities in the environment and adapts to an uncertainty associated with those regularities are central questions in cognitive and behavioral neuroscience. Here, we used multiple metrics based on information theory to investigate the effects of different types of uncertainty on learning and choice behavior. We found that rodents and primates differed in how they adapted to uncertainty in the environment. Mice employed option-dependent strategies more often, after slowly identifying the better option, and chose to stay on the better rewarding option to exploit their current estimates of reward probabilities. In contrast, monkeys employed reward-dependent strategies more often while switching between choice options more frequently. Fit of choice data using RL models showed that this difference in strategy was mediated via higher learning rates in monkeys, consistent with the interpretation that such strategy requires stronger weighting of immediate reward outcomes. In addition, the inverse temperature representing choice consistency was higher in mice, as they consistently stayed on the better option more than monkeys.

Some of the similarities and differences between mice and monkeys can be explained as below. First, mice exhibited an overall more consistent choice strategy, indicated by the lower values of H(str), ERDS, and EODS than those of monkeys. The higher performance (P(Better)) in mice suggests that the lower entropy was mainly due to these animals staying on the better option more consistently than monkeys. We note that this difference could be in part derived from slight differences in tasks rather than reflecting some intrinsic intra-species differences. In the mouse experiment, the same reward schedule was used throughout each of the entire session, while the reward probabilities reversed between two sides at every block switch (**Figure 2C**). In contrast, in the monkey task, reward schedules were randomly assigned at every block, making it more challenging to determine the better option (**Figure 2D**). Furthermore, mice on average experienced more than twice the number of block reversals than that of monkeys during each session of the experiment (**Methods**). These two characteristics of the mouse experiment (use of only two reward probabilities in each session and more frequent block switches within a session) could facilitate the distinction between the better and worse options, resulting in more adoption of an option-based strategy. In addition, reward delivery in the mouse experiment was from two separate (left and right) ports instead of the one spout used in the monkey experiment. All of these differences could make it easier for mice to adopt an option-based strategy.

Furthermore, the difference in the reward rates between two options (Δ*P*) was smaller in the monkey experiment, suggesting that the task difficulty was greater for monkeys. Interestingly, we found some evidence that within each species, increased task difficulty in terms of reward probability was accompanied by the animals adopting a more reward-based strategy (**Figure S9**). Therefore, higher task difficulty in the monkey experiment could promote a reward-based strategy and lead to a more switching behavior. Specifically, the monkeys showed a short-term tendency to switch between options and a longer-term tendency to repeat choices for the same options—a pattern that can be revealed by inclusion of a choice-history variable in a logistic regression on the monkeys’ choices (Grabenhorst et al., 2019) and that has previously been observed in monkeys performing matching tasks (Lau & Glimcher, 2005). The monkeys’ higher tendency to switch may have also resulted from the absence of a “changeover delay,” a manipulation sometimes used to encourage matching behavior, whereby reward would only be delivered on the second trial following a switch between options (Sugrue et al., 2004). Interestingly, H(str) was positively correlated with performance P(Better), indicating that a more equal mix of stay and switch facilitated winning of rewards in monkeys. In contrast, H(str) was negatively correlated with performance in mice, as they primarily employed a staying strategy to exploit the better rewarding option. Future empirical work should test the influence of task conditions on these observed species differences.

The overall difference between mice’s and monkeys’ adoption of stay vs. switch strategies could also explain how they adjusted to uncertainty. More specifically, the greater uncertainty in the reward environment (measured by Δ*P*) had different effects on mice and monkeys. Mice adjusted their strategies by incorporating switching strategies more often, which resulted in increased randomness in strategies reflected by higher entropies (H(str), ERDS, and EODS). Monkeys similarly increased their switching behavior, but even more so than their baseline strategy (which was already very random) such that the overall H(str) was reduced, signaling higher consistency.

The difference in the baseline entropy of strategy (H(str)) between mice and monkeys also partly accounts for the observed difference in their adjustment to expected block switches. The exploitation strategy characterizing the mice behavior—consistently staying on the more rewarding option—is a form of option-dependent strategy which requires a clear sense or confidence about what the better option is. Therefore, in the anticipation or perception of block switches, the link between previous choice option and the subsequent action would be expected to decline as the animal begins to explore the alternative option. This is what the MIOS metric in mice revealed (comparison of before/after metrics in **Figure S7** or **Figure S12I**). In contrast, monkeys primarily depended on reward-dependent strategies. Accordingly, in the event of expected block switches, the link between reward feedback and the subsequent action was most significantly reduced (MIRS in **Figure S7E** and **Figure S12F**), as winning a reward did not lead to staying behavior as consistently as before.

Here, we used the difference between the baseline probability of reward for the better and worse options to quantify expected uncertainty in the environment. This was based on the notion that a larger difference in reward probabilities is associated with less uncertainty in identifying the better option using reward feedback. Alternatively, one could use the variance in objective or subjective reward probability to measure risk in an economic sense, as such “variance risk” can drive choice behavior and neural activity differently than subjective value (Schultz et al., 2008; Grabenhorst, et al., 2019). In this study, we used uncertainty because in both tasks, reward probabilities are unknown and must be learned from reward feedback (De Groot & Thurik, 2018; Soltani & Izquierdo, 2019) as opposed to their distributions being fully accessible to the subject (Farashahi et al., 2018; Woo et al., 2022). We examined how such uncertainty influences choice behavior and learning.

Comparing the effects of reward uncertainty (ΔP) and block switch expectation, we found that the effects of block switch expectation were more subtle compared to that of Δ*P*. That is, for all of our analyses comparing the extent of these two effects (**Figure 4, 5**; **Figure S11, S12**), reward uncertainty had a greater impact on behavior than block switch expectation for both species. This is expected because anticipation of block switches requires longer-term statistics of reward outcomes compared to those related to determining the better option, as shown in a recent study (Atilgan et al., 2022). The animals could, for example, return to their previous strategy after enough reward feedback has signaled that actual block switch has not occurred yet. Still, the entropy-based measures captured such effects from LE blocks, by applying the assumption that expectation of block switches can have more clear effects toward the end of blocks. This also demonstrates the utility of the entropy metrics, which are capable of detecting nuanced shifts in choice behavior if measured from an appropriate binning of trials.

Finally, the observation that expectation of block switches depends on the level of uncertainty suggests interdependence between different types of uncertainty (Soltani & Izquiredo, 2019; Piray & Daw, 2021). In a recent study, Piray & Daw (2021) provided a normative account that uncertainty (stochasticity) reduces the learning rate, whereas volatility increases it. Regarding uncertainty, we similarly observed lower mean learning rates for more uncertain conditions. However, on the influence of volatility, previous studies have shown mixed results (Behrens et al., 2007; Donahue and Lee, 2015; Farashahi et al., 2019). Consistent with previous work (Donahue and Lee, 2015; Farashahi et al., 2019), we found decreased rather than increased learning rates in both species. Moreover, we found that the learning rates were not influenced by the mediating effects of greater uncertainty on the block switch expectation.

These results raise a possibility that, while learning rates could be implicated in adjustment to volatile environments, the effect of expectation arising from observation of long-term regularities in the environment might more directly involve other cognitive mechanisms. For example, it has been suggested that active representations of alternative models about the environment (task set) are stored in long-term memory to be retrieved when the corresponding context reoccurs (Collin & Koechlin, 2012; Soltani & Koechlin, 2022). The neural system for this retrieval likely involves extensive interaction between the prefrontal cortex and the hippocampus, the latter proposed to be involved in the representation of unexpected uncertainty (Payzan-LeNestour et al., 2013) and one-shot learning (Lee et al., 2015) in humans. Further studies in rodents and primates could directly probe the causal role for the hippocampus in learning under different forms of uncertainty.

There are limitations and additional considerations related to our findings. Foraging in the wild consists of exposure to time-delays and spatial separation between options, so more naturalistic dynamic foraging tasks could be developed to increase parametric space such that a wider range of values could be explored (Constantino & Daw, 2015; Hall-McMaster et al., 2021). Indeed, the longer time horizon and expansive range size to forage in nonhuman primates compared to rodents influence how the species predict, evaluate, and act on rewards; mediated by a more specialized and expansive prefrontal cortex in the primate brain (Wittmann et al., 2016; Rudebeck & Izquierdo, 2022). Additionally, though we found that stochastic behavior and corresponding entropy metrics were affected by uncertainty of the better option in both mice and monkeys, these effects may be mediated by different neural mechanisms given that mice responded by actions, whereas monkeys responded to stimuli. Previous work has shown that (freely moving) rodents tend to have greater difficulty learning stimulus-based probabilistic outcomes compared to spatial-based ones (Harris et al., 2021; Chen et al., 2021), whereas monkeys do well in both modalities (Costa et al., 2016; Rothenhoefer et al., 2017; Taswell et al., 2021). Consistently, different neurons in the dorsolateral prefrontal cortex of monkeys have been shown to track reward uncertainty associated with both stimuli and actions (Grabenhorst et al., 2019). This points to a need to disambiguate the rodents’ natural tendency to navigate and learn spatially, versus the influence of task conditions that enhance stable choice such as in this dynamic foraging task.

## Conclusion

Together, our results illustrate how higher-order statistics of the environment influence learning and choice strategies. We show that entropy-based metrics can be useful for investigating multiple aspects of adaptive behavior under different forms of uncertainty. By introducing the notion of block switch expectation, we showed that shorter than expected blocks and longer than expected blocks were marked by quantitatively different consistencies in behavior as captured by the entropy-based metrics. These findings suggest that mice and monkeys can use reward history to form long-term expectations about the environment and adjust their behavior accordingly. Such expectations of block switches were characterized by increased randomness in choice strategy, similar in its quantity to those induced by the actual block switches. Therefore, our study demonstrates how model-free (entropy-based metrics) and model-dependent (RL models) methods can complement each other to elucidate behavioral adjustments and underlying mechanisms with finer details.

## Supplementary Information

The online version contains supplementary material available at XXX.

## Acknowledgements

We would like to thank Ethan Trepka for sharing the codes for computation of entropy-based metrics, Dmitriy Lisitsyn for suggestions about the permutation test and bootstrapping method, and Jane Han for helpful suggestions about figure aesthetics.

## Declarations

### Authors’ contribution

*Conceptualization of the study:* JHW, AS; *Mouse experiment conceptualization & data collection:* BAB, JYC; *Monkey experiment conceptualization & data collection:* KT, FG, WS; *Analyses of data and illustration:* JHW, CGA, AI, AS; *Implementation of algorithms, models, analyses, and plots:* JHW; *Original draft:* JHW, CGA, AI, AS; *Review & editing:* all authors.

### Availability of codes, data & materials

The codes used for analysis and plotting the figures are available online at https://github.com/SoltaniLab. The data presented are available upon reasonable request from the authors. The reported experiments were not preregistered.

## Supplementary Materials

**Figure S1.**
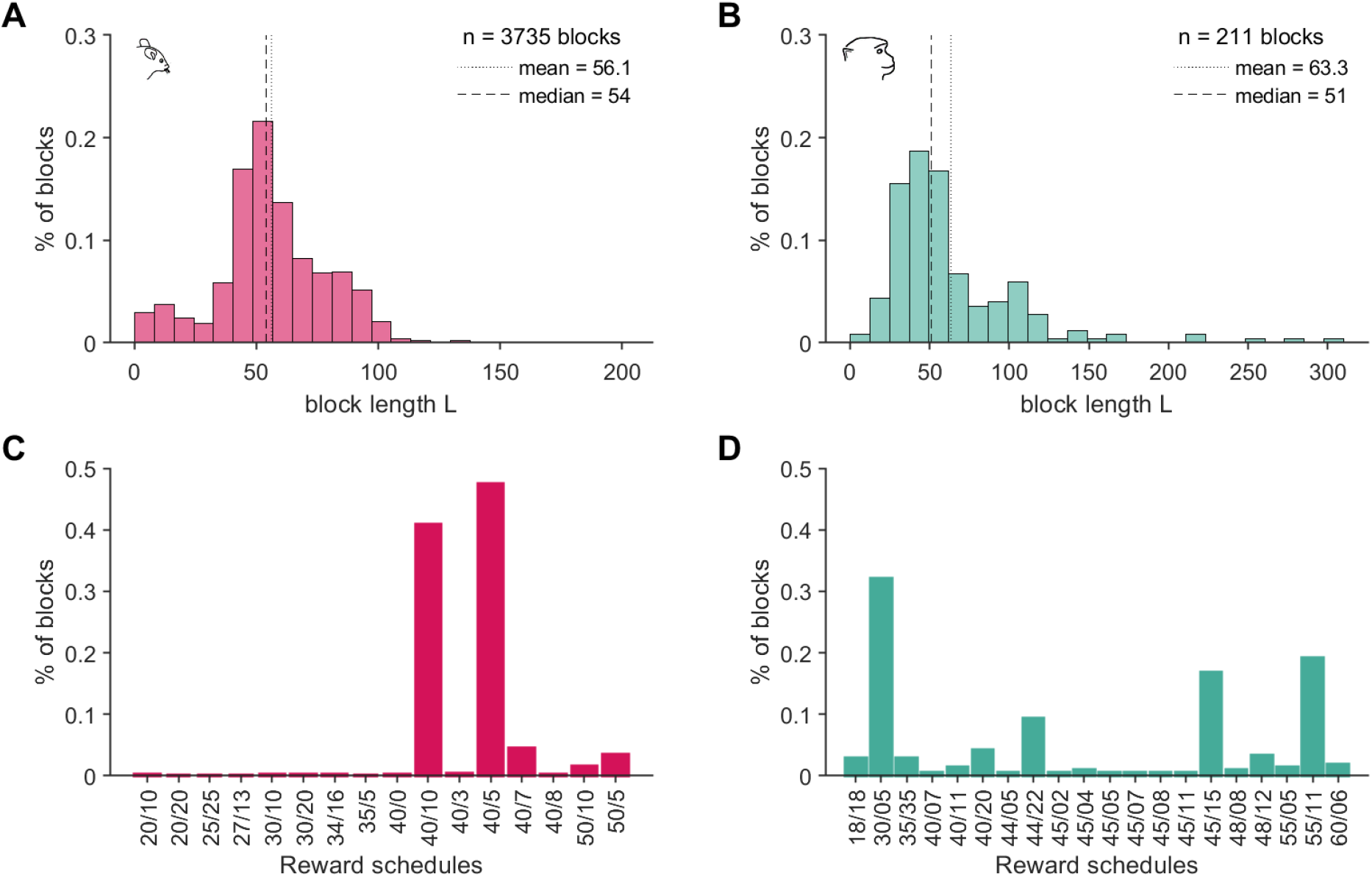
Distributions of block lengths and reward probabilities in dynamic foraging tasks in mice and monkeys. Plots show distributions of block lengths and reward schedules in the mouse (A, C) and monkey (B, D) experiments. The two numbers in the reward schedule correspond to reward probabilities (in percentage) of better and worse options (defined based on location for mice and visual stimuli for monkeys) in each block. For some blocks, the two choice options had the same reward probabilities (e.g., 35/35). Total *n* = 18 mice performed on average 209.3 blocks per each subject. Total *n* = 2 monkeys performed on average 125.5 blocks (first monkey: 182 blocks, second monkey: 69 blocks). The majority of the blocks (~89%) in mice followed either a 40/10 or a 40/5 schedule. The majority of the blocks (~70%) in monkeys followed a 30/5, 45/15, or a 55/11 schedule.

**Figure S2.**
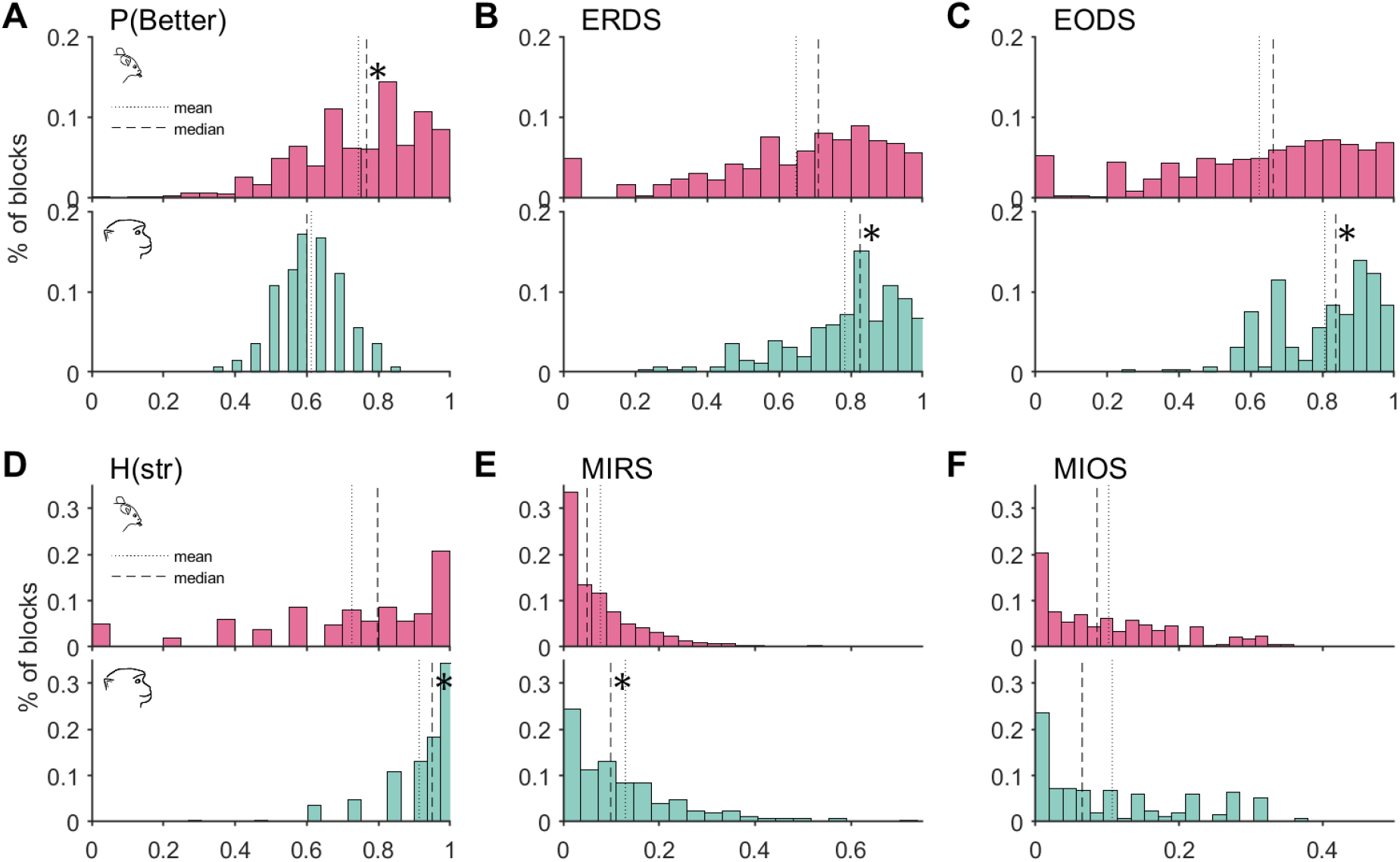
Distributions of main behavioral metrics in mice and monkeys. Plotted are distributions of the behavioral metrics in mouse (top panel, red) and monkey (bottom panel, teal) experiments, including performance or probability of choosing a better option (A), entropy of reward-dependent strategy (B), entropy of option-dependent strategy (C), baseline entropy of strategy (D), mutual information of reward and strategy (E), and mutual information of previous choice option and strategy (F). Asterisk indicates the distribution with significantly larger CDF from intra-species comparison, determined by Kolmogorov-Smirnov test (*p* < .05). All metrics have minimum values of 0 and maximum values of 1. Due to log-transformation in the computation of entropy-based metrics, most of the metrics are highly skewed. To avoid confounds with the data size and the computed entropy values, we kept the same bin size (number of trials) from each block for computing the metrics. For mouse data, metrics are computed from the last *N* = 30 trials in the block. For monkey data, *N* = 20. This was chosen to minimize the number of excluded blocks while keeping its proportion consistent across datasets (see **Methods**).

**Figure S3.**
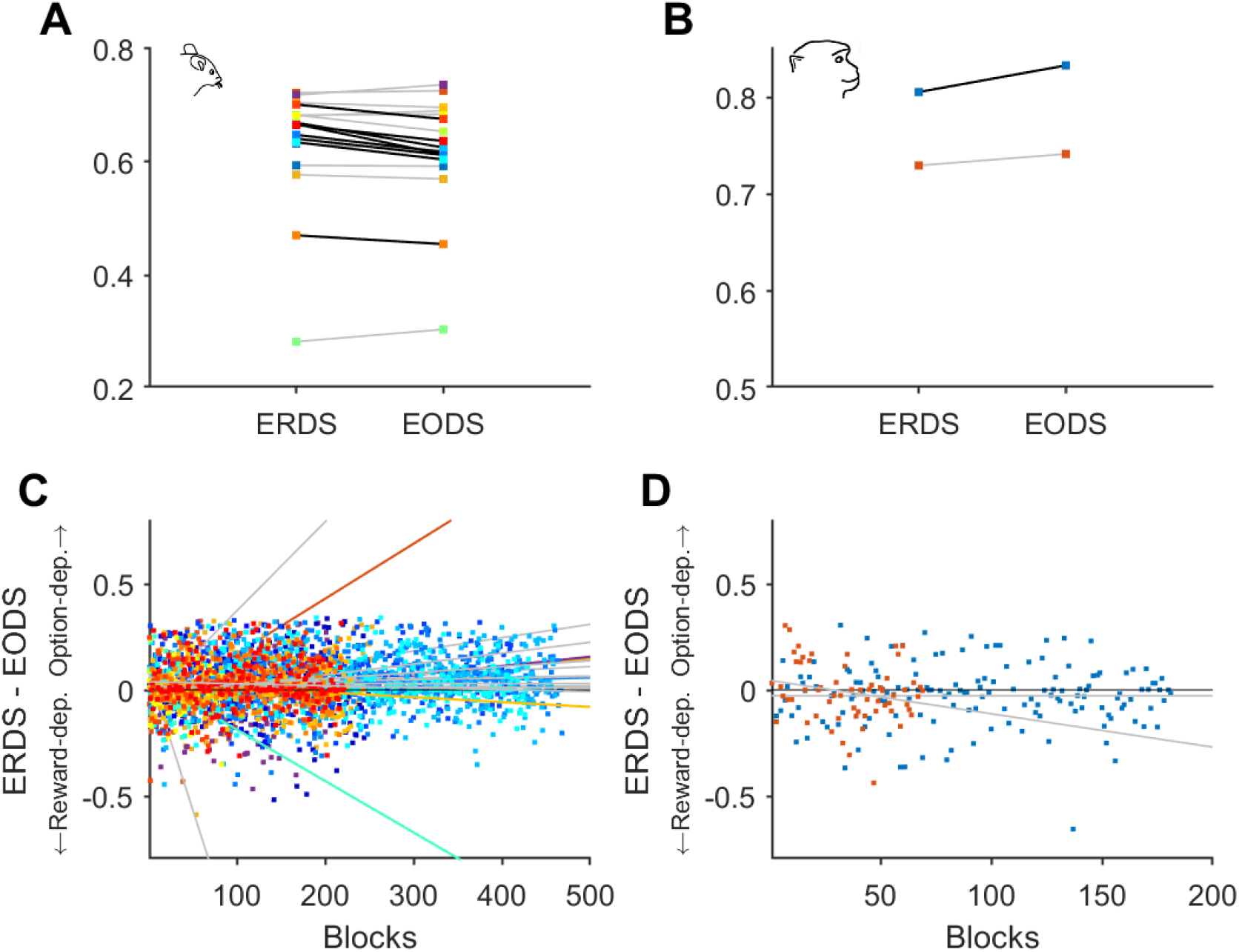
Adopted strategy is consistent across individuals within each species. (**A–B**) Mean ERDS and EODS for individual mice (A) and monkeys (B) is shown with different colors. Significant difference between two values (using two-sided paired-sample t-test) is indicated with a black line connecting data for each individual. Overall, thirteen mice (72% of total) had lower EODS than ERDS, indicating preference for an option-dependent strategy. One monkey showed significantly lower ERDS than EODS (*p* = .0206), whereas the other monkey followed a similar trend (*p* = .4980). (**C–D**) Plots show the difference in ERDS and EODS for individual mice (C) and monkeys (D) as a function of the number of blocks performed. Individuals with significant slopes are shown with colored lines. Majority of mice tended to prefer an option-dependent strategy more strongly as time passed, whereas the two monkeys adopted a more reward-based strategy with time.

**Figure S4.**
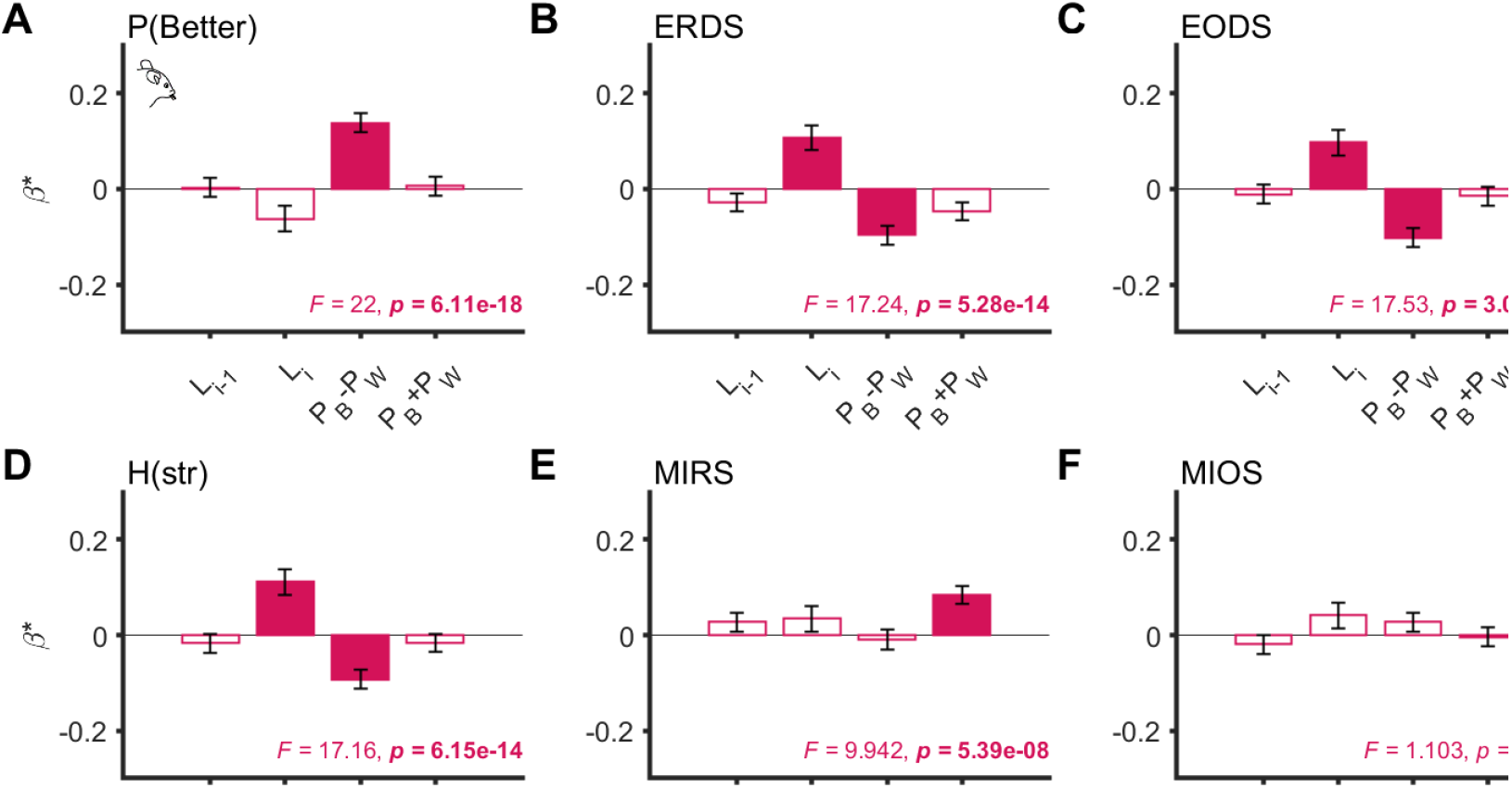
Multiple regression on mice’s behavioral metrics. Bar plots show the standardized coefficients of regressors, corresponding to task variables representing different types of uncertainty. The bottom of each panel reports the F-statistic and its p-value of the regression model. Significant predictors are indicated with a filled color bar. *L_i-1_* = previous block length. *L_i_* = length of the current *i*-th block. *P_B_* - *P_W_* = difference between reward probabilities of better (*P_B_*) and worse (*P_W_*) options (see Methods). *P_B_* + *P_W_* = sum of the two reward probabilities. All behavioral metrics were computed from the last 30 trials of each block, to avoid confounds with data size and computed entropy values by ensuring the same bin size for every block. (**A**) Response variable in the regression is performance (probability of choosing the better option during each block, P(Better)). Larger *P_B_* - *P_W_* or less expected uncertainty predicted better performance (***β***\* = 0.156, *p* = 3.32×10^-15^). Longer *L_i_* was associated with worse performance (***β***\* = −0.0539, *p* = .0416) but the significance did not survive after correcting for multiple comparisons of four (number of independent variables). (**B)** Response variable: entropy of reward-dependent strategy (ERDS), equal to *H*(*str|rew*). Higher ERDS value indicates a more random response strategy on reward feedback. Longer *L_i_* and smaller *P_B_* - *P_W_* predicted more random reward-dependent strategy (*L_i_*: ***β***\* = 0.107, *p* = 5.73×10^-5^; *P_B_* - *P_W_*: ***β***\* = −0.096, *p* = 1.50×10^-6^) (**C**) Response variable: entropy of option-dependent strategy (EODS), equal to *H*(*str|opt*). Higher EODS value indicates a more random response strategy on the previous chosen option. Similar to ERDS, Longer *L_i_* and smaller *P_B_* - *P_W_* predicted more random option-dependent strategy (*L_i_*: ***β***\* = 0.097, *p* = 2.54×10^-4^; *P_B_* - *P_W_*: ***β***\* = −0.103, *p* = 2.90×10^-7^). (**D**) Response variable: entropy of strategy, H(*str*). This quantity represents the baseline entropy of stay or switch strategy, independent of any information from the previous trial. Longer *L_i_* and smaller *P_B_* - *P_W_* predicted more random strategy (*L_i_*: ***β***\* = 0111, *p* = 2.91×10^-5^; *P_B_* - *P_W_*: ***β***\* = −0.093, *p* = 3.28×10^-6^). (**E**) Response variable: mutual information of reward and strategy (MIRS), equal to I(*rew*; *str*). Note *ERDS* = H(*str*) - MIRS. Larger *P_B_* - *P_W_* predicted more shared information between reward and strategy (***β***\* = 0.084, *p* = 2.09×10^-5^). (**F**) Response variable: mutual information of option and strategy (MIOS), equal to *I*(*opt; str*). Note EODS = H(*str*) - MIOS. No predictors were significant.

**Figure S5.**
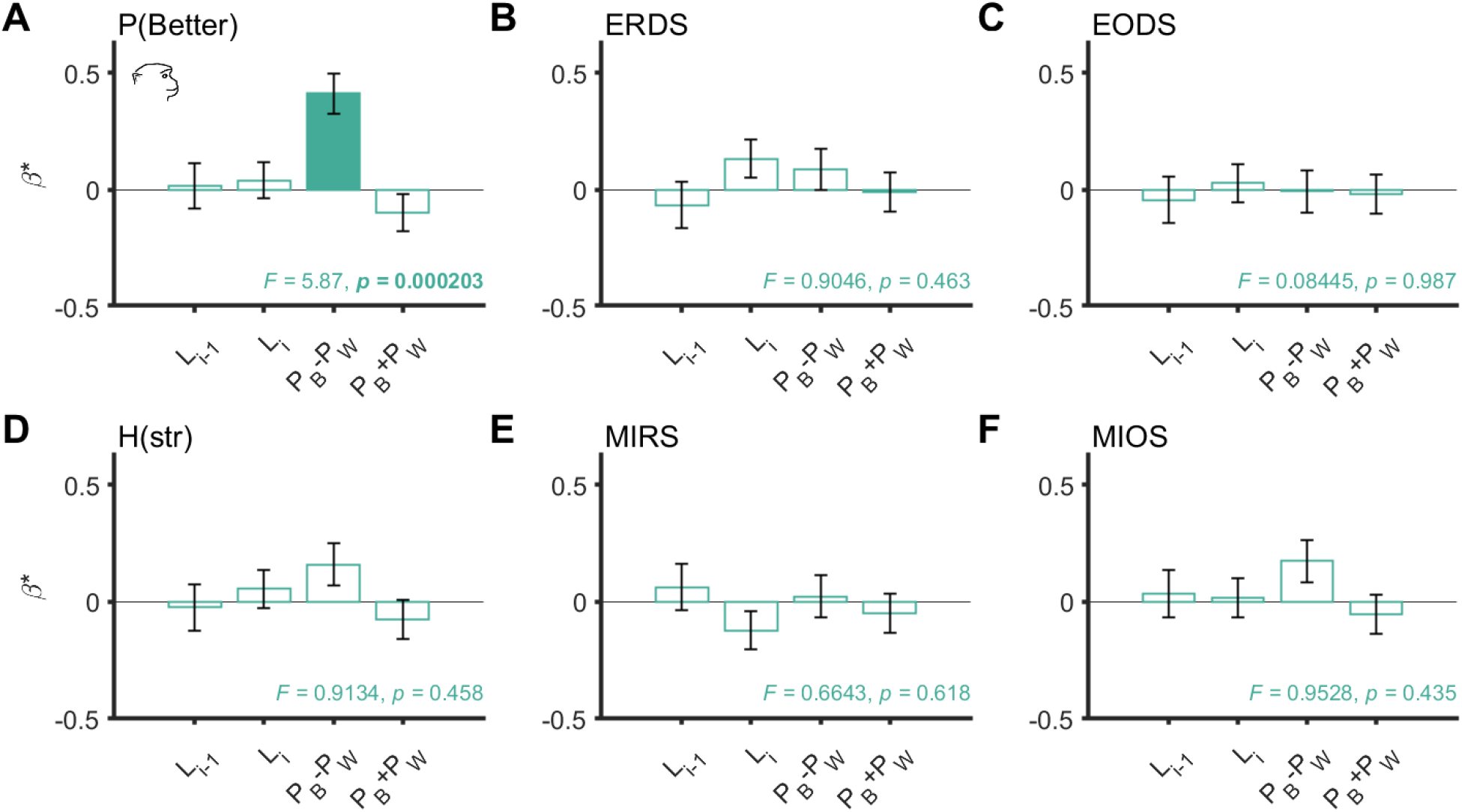
Multiple regression on monkeys’ behavioral metrics. Bar plots show the standardized coefficients of regressors, corresponding to task variables representing different types of uncertainty. The bottom of each panel reports the F-statistic and its p-value of the regression model. Significant predictors are indicated with a filled color bar. *L_i-1_* = previous block length. *L_i_* = length of the current block *i*. *P_B_* - *P_W_* = difference between reward probabilities of better (*P_B_*) and worse (*P_W_*) options (see Methods). *P_B_* + *P_W_* = sum of the two reward probabilities. All behavioral metrics were computed from the last 20 trials of each block, to avoid confounds with data size and computed entropy values by ensuring the same bin size for every block. (**A**) Response variable in the regression is performance or probability of choosing the better option during each block, P(Better). Larger *P_B_* - *P_W_* or less uncertainty predicted better performance (***β***\* = 0.412, *p* = 3.76×10^-6^). (**B**) Response variable: entropy of reward-dependent strategy (ERDS), equal to *H*(*str*|*rew*). Longer *L_i_* was associated with more random reward-dependent strategy but not significantly (***β***\* = 0.133, *p* = .106). (**C**) Entropy of option-dependent strategy (EODS), equal to *H*(*str*|*opt*). (**D**) Entropy of strategy, H(*str*). Larger *P_B_* - *P_W_* or less uncertainty was associated with more random strategy but not significantly (***β***\* = 0.159, *p* = .075). (**E**) Mutual information of reward and strategy (MIRS), equal to I(*rew*; *str*). Note *ERDS* = *H*(*str*) - MIRS. Longer *L_i_* was associated with less shared information but not significantly (***β***\* = −0.122, *p* = .140). (**F**) Mutual information of option and strategy (MIOS), equal to I(*opt*; *str*). Note *EODS* = H(*str*) - MIOS. Larger *P_B_* - *P_W_* or less uncertainty was associated with more random option-dependent strategy but not significantly (***β***\* = 0.173, *p* = .057).

**Figure S6.**
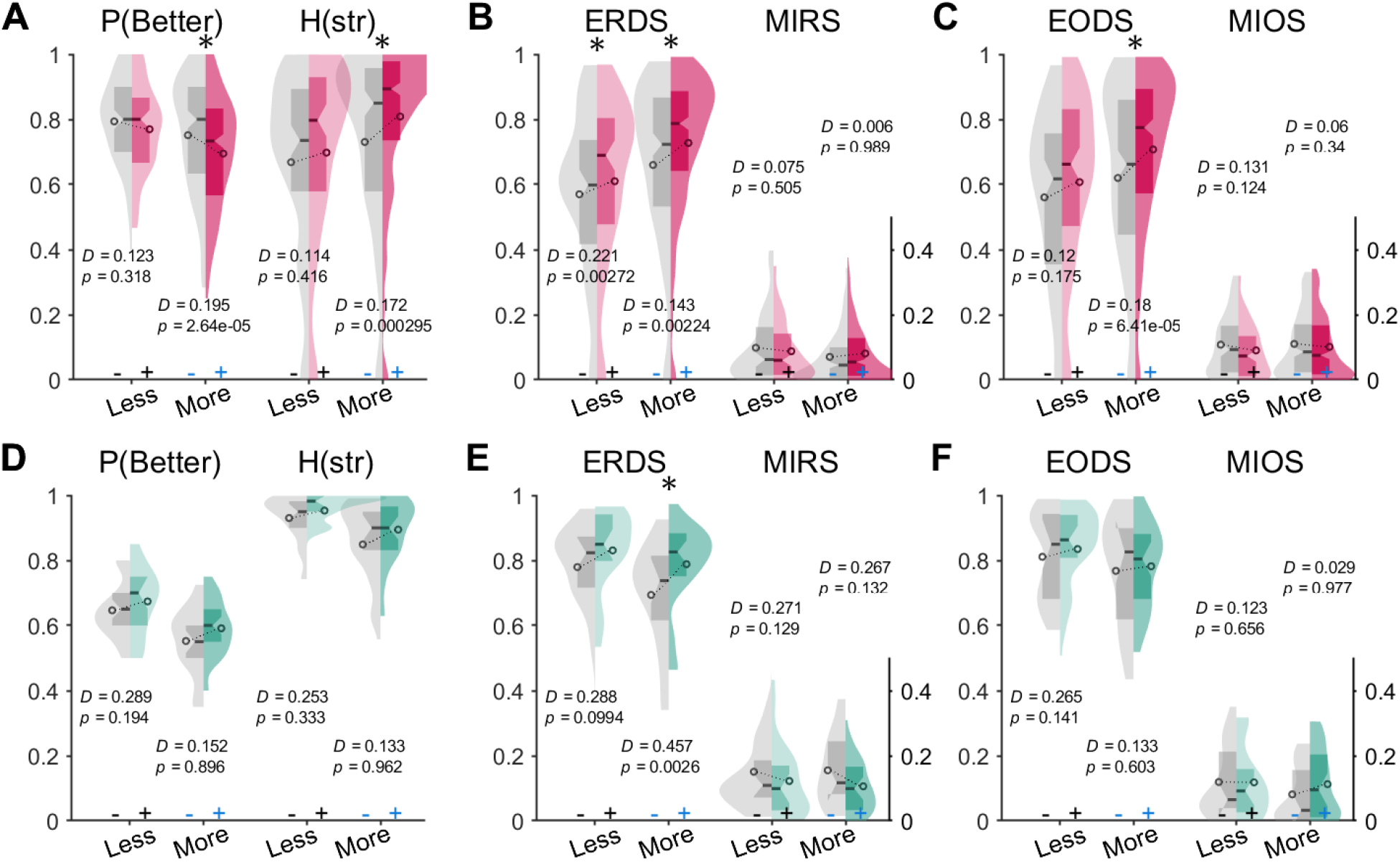
Expectation of block switches depends on the level of uncertainty. Plotted are distributions showing combined effects of uncertainty and block switch expectation on choice behavior for mice (top, **A–C**) and monkeys (bottom, **D–F**). For each metric, blocks are divided into four categories: SE and less uncertain (gray, *Less*-), LE and less uncertain (colored, *Less*+), SE and more uncertain (gray, *More*-), and LE and more uncertain (colored, *More+*). Asterisks show results from a two-sample Kolmogorov-Smirnov test. All tests are two-sided, corrected for multiple comparisons of four. (**A**) In mice, performance P(Better) was significantly decreased for LE blocks in a more uncertain environment only. Similarly, H(str) or randomness in strategy was significantly increased for LE blocks in a more uncertain environment only. (**B**) In contrast, ERDS was larger for LE blocks in both less and more uncertain environments. (**C**) The effect of block switch expectation on EODS was significant in more uncertain environment only. (**D–F**) For monkeys, the only significant difference between SE and LE was found for ERDS in a more uncertain environment.

**Figure S7.**
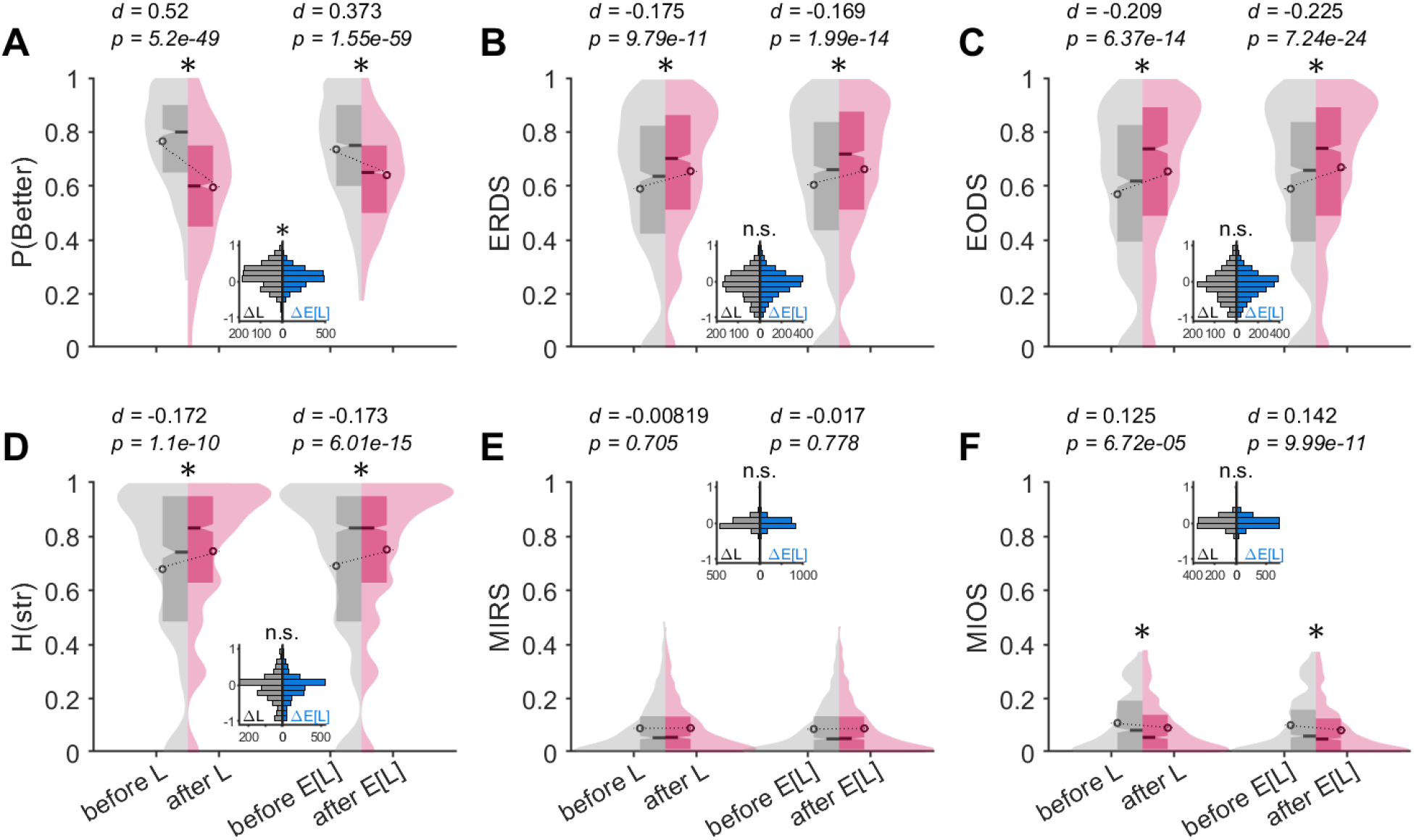
Actual and expected block switches have similar effects on mouse choice behavior. Plotted are distributions of metrics computed from 20 trials around actual block switches (*L* on the left, containing SE blocks only) and expected block switches (*E[L]* on the right, containing LE blocks only). Reported values are Cohen’s *d* and p-values based on paired-sample t-test comparing before and after actual and expected block switches (one-sided t-test, with a priori hypothesis that entropy increases and P(Better) (mutual information) decreases after actual and expected block switches). Insets show distributions of paired differences around actual (ΔL) and expected (ΔE[L]) block switches, with asterisks indicating significant difference between two means based on independent-sample t-test (two-sided; n.s. = not significant, *p* > .05). (**A**) During 20 trials after the expected block switches point (for LE blocks, right), performance significantly decreased, similar to the behavior after the actual block switches (for SE blocks, left). (**B, C**) Conditional entropies, ERDS and EODS, similarly increased after both actual and expected block switches. (**D**) Baseline entropy of strategy, H(str), similarly increased after both actual and expected block switches. (**E**) MIRS were unaffected by both actual and expected block switches. (**F**) MIOS significantly decreased after both actual (*L*, Cohen’s *d* = 0.125, *p* = 6.72×10^-5^) and expected (*E*[*L*], Cohen’s *d* = 0.142, *p* = 9.99×10^-11^) block switches, indicating a change in the option-dependent strategy.

**Figure S8.**
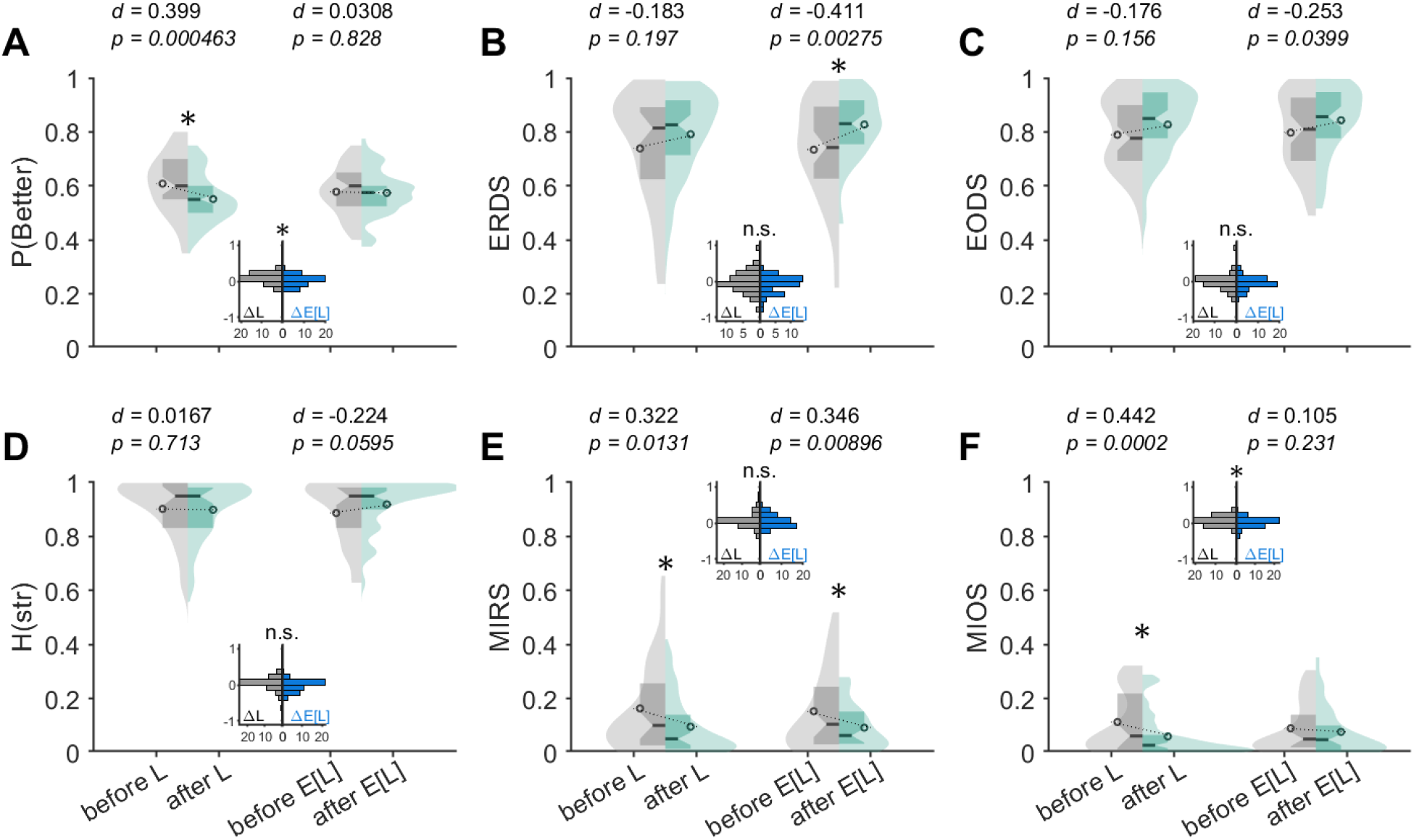
Actual and expected block switches have similar effects on monkey choice behavior. Plotted are distributions of metrics computed from 20 trials around actual block switches (*L* on the left, containing SE blocks only) and expected block switches (*E*[*L*] on right, containing LE blocks only). Reported values are Cohen’s *d* and p-values based on paired-sample t-test comparing before and after actual and expected block switches (one-sided t-test, with a priori hypothesis that entropy increases and P(Better) and mutual information decrease after actual and expected block switches). Insets show distributions of paired differences around actual (ΔL) and expected (ΔE[L]) block switches, with asterisks indicating significant difference between two means based on independent-sample t-test (two-sided; n.s. = not significant, *p* > .05). (**A**) Performance decreased after actual block switches (left), but not significantly after expected block switches (right). (**B**) ERDS increased after both actual and expected block switches. The increase after actual block switches (left) was moderate but not statistically significant (Cohen’s *d* = −0.183, *p* = .197). (**C, D**) EODS and H(str) were unaffected by both actual and expected block switches. (**E**) MIRS significantly decreased after both actual (*L*, Cohen’s *d* = 0.322, *p* = .0131) and expected (*E*[*L*], Cohen’s *d* = 0.346, *p* = .00896) block switches, indicating a change in the reward-dependent strategy. (**F**) MIOS significantly decreased after actual but not expected block switches.

**Figure S9.**
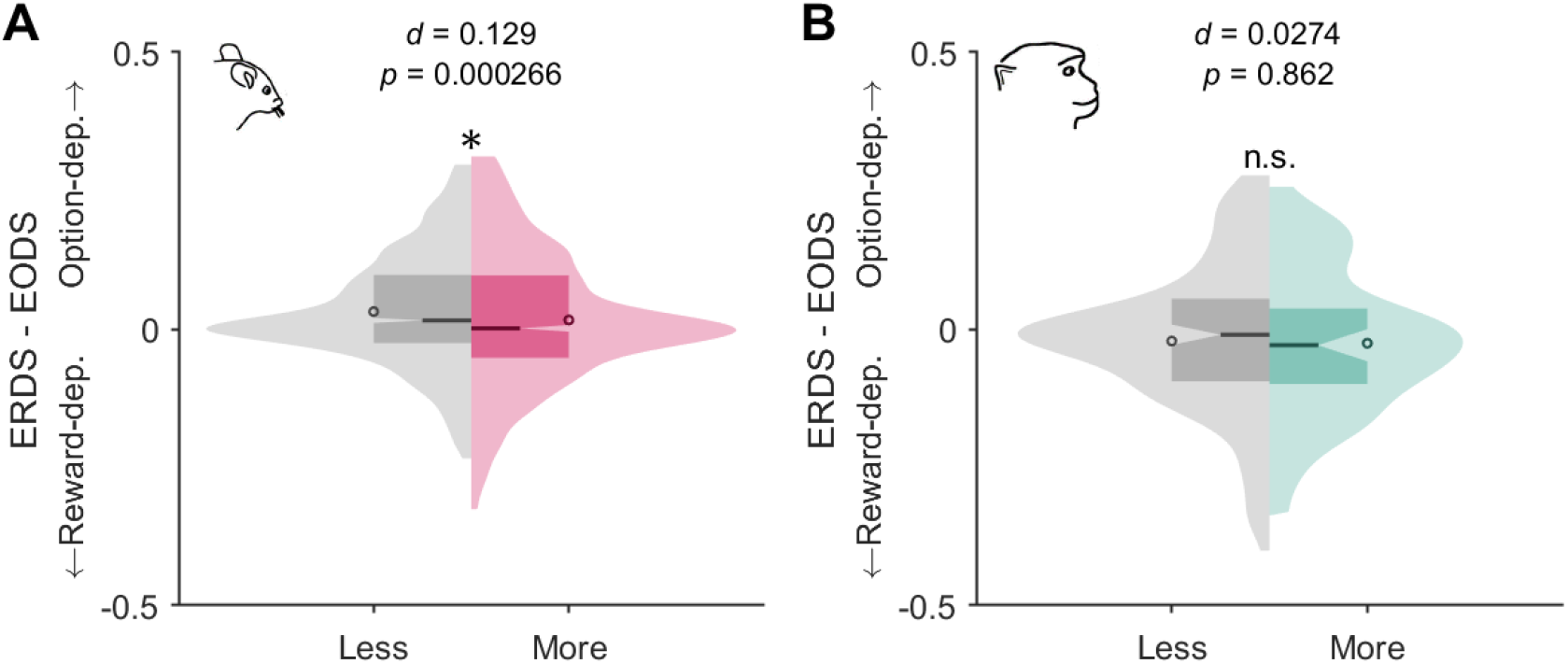
Preferred strategy depends on the level of uncertainty in determining the better option. Plotted are distributions of the difference in ERDS and EODS in less or more uncertain environments separately for mice (A) and monkeys (B). Reported are Cohen’s *d* and p-values based on two-sample t-test, and the asterisk indicates a statistically significant difference. Mice exhibited significantly a more reward-dependent strategy in more uncertain environments, suggested by a decrease in (ERDS - EODS). Monkeys tended toward a more reward-dependent strategy in more uncertain environments.

**Figure S10.**
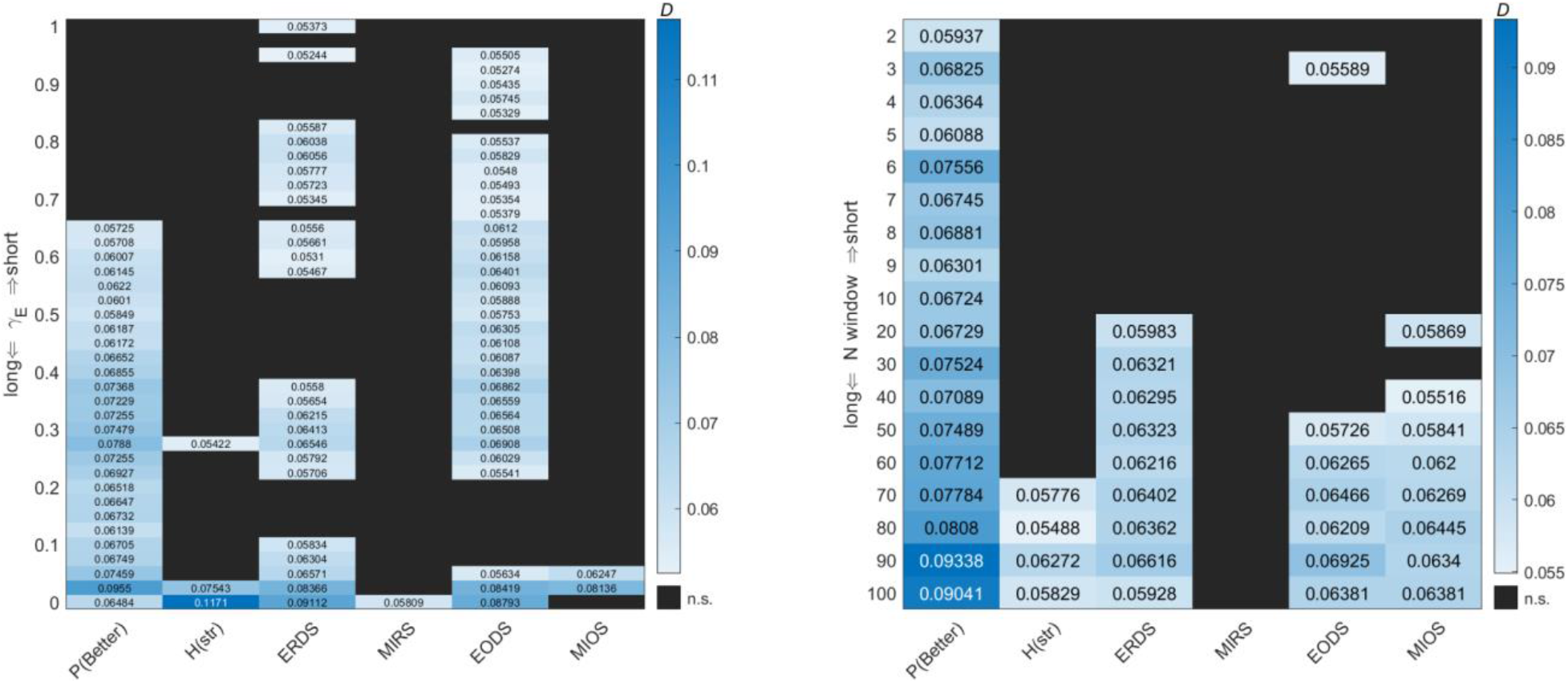
Comparison of different timescales for integration of expectation of block length in mice. Reported are the *D* statistics of the two-sample Kolmogorov-Smirnov test, with a null hypothesis that the two samples of metrics (from SE and LE blocks) are from the same continuous distribution (two-sided test). n.s. = not significant (*p* > .025). Left panel: *D* statistics as a function of *γ_E_* parameter used to compute expected block length *E*[*L*] (Eq. 7 in Methods). Larger *γ_E_* values correspond to shorter timescales and smaller values to longer timescales for integration of expected block length. Right panel: *D* statistics as a function of window size *N* used to compute expected block length *E*[*L*]_*N*_ (Eq. 9 in Methods). Smaller *N* values correspond to shorter timescales, and smaller values to longer timescales for integration of expected block length. For mice, the distinction between SE and LE blocks tended to be more pronounced for longer timescales of *E*[*L*].

**Figure S11.**
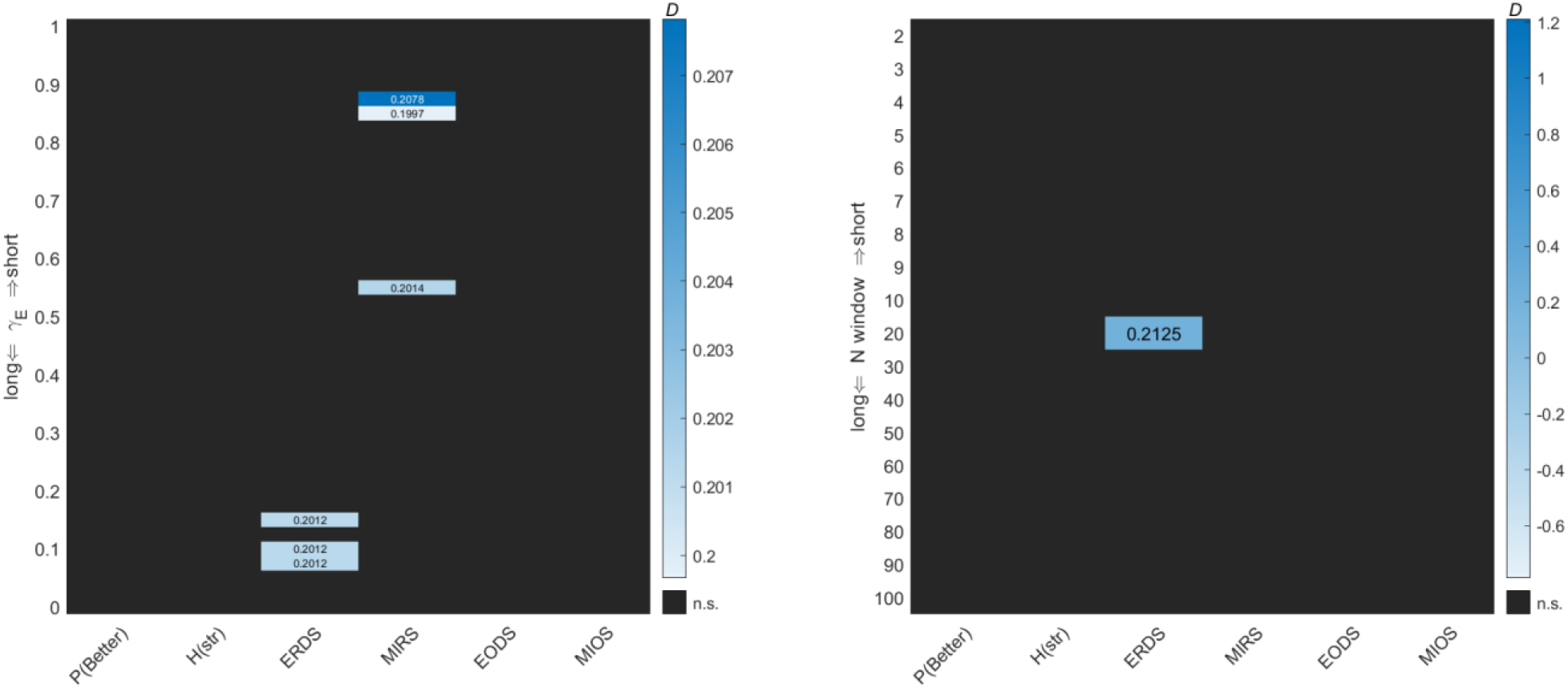
Comparison of different timescales for integration of expectation of block length in monkeys. Reported are the *D* statistics of the two-sample Kolmogorov-Smirnov test, with a null hypothesis that the two samples of metrics (from SE and LE blocks) are from the same continuous distribution (two-sided test); n.s. = not significant (*p* > .025). Left panel: *D* statistics as a function of *γ_E_* used to compute expected block length *E*[*L*] (Eq. 7 in Methods). Larger *γ_E_* values correspond to shorter timescales and smaller values to longer timescales for integration of expected block length. Right panel: *D* statistics as a function of window size *N* used to compute expected block length *E*[*L*]_*N*_ (Eq. 9 in Methods). Smaller *N* values correspond to shorter timescales, and smaller values to longer timescales for integration of expected block length. For monkeys, the distinction between ERDS among SE and LE blocks tended to be significant only for relatively longer timescales of *E*[*L*] (*γ_E_* = ~0.1 and *N* = 20).

**Figure S12.**
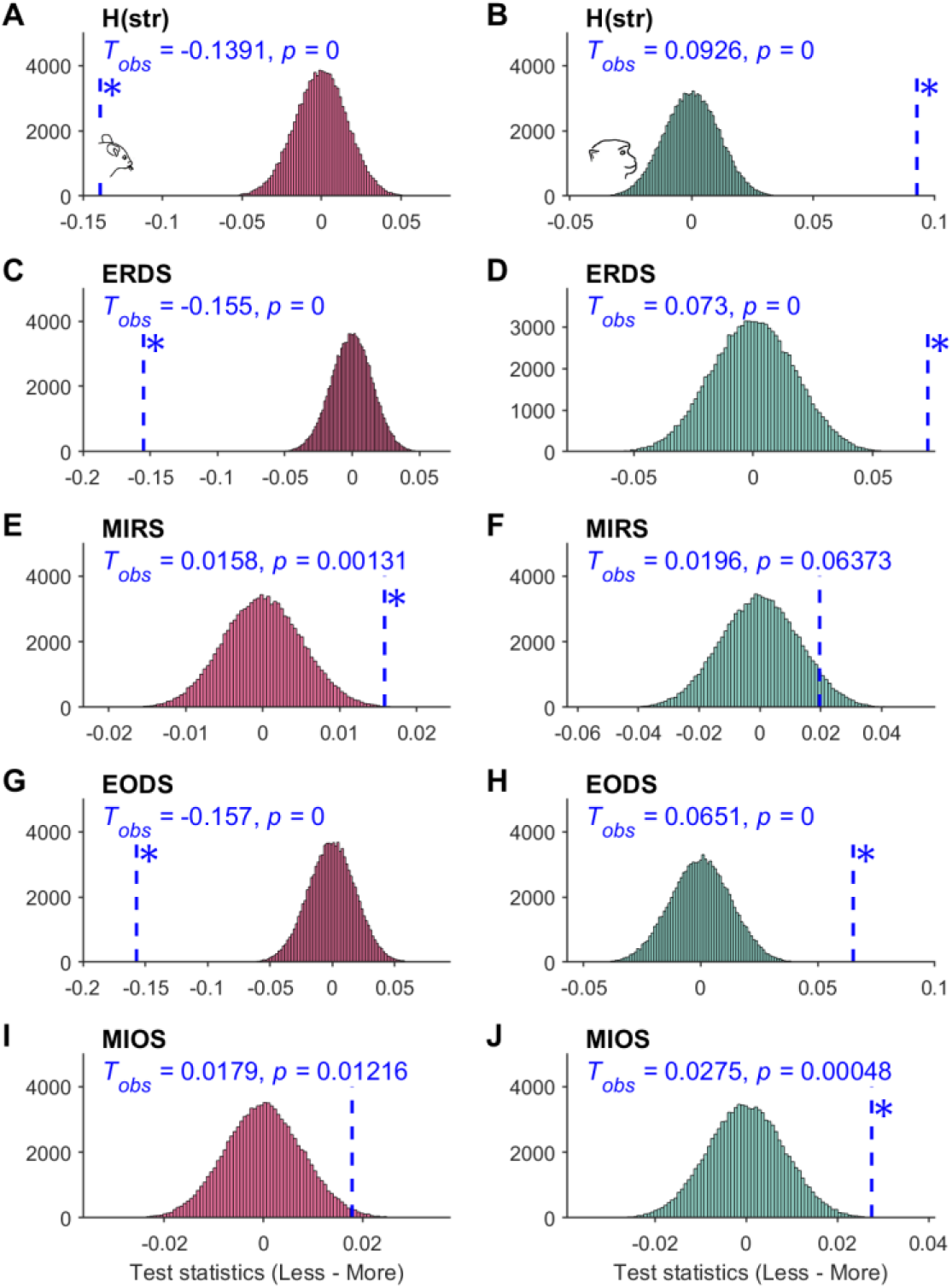
Effects of reward uncertainty on mouse and monkey behavior shown by permutation test. Plots show distributions of test statistics from *N* = 100,000 sampled permutations in mice (column 1) and monkeys (column 2), under the null hypothesis that less and more uncertain conditions have the same entropy measures. Vertical dotted lines in blue indicate the actual observed test statistics (T_obs_), which is the difference between a given entropy metric for the two block types. P-values indicate the probability of observing test statistics as extreme as the observed value (one-tailed). Asterisks indicate *p* < 0.01. For each permutation, we randomly assign blocks into two groups with sample size *n_1_* or *n_2_*, corresponding to the total number of blocks in *less* or *more uncertain* conditions, respectively. A single entropy measure is computed for each type of block by concatenating all trial vectors from the sampled permutation. Test statistic for each sample is then computed by taking the difference in metrics between the first and the second group. (**A,C,E,G,I**) Mouse data in red. More uncertain blocks had significantly larger H(str), ERDS, and EODS, indicating an overall increase in randomness of choice strategy. MIRS was also significantly smaller, indicating a decreased link between reward outcome and subsequent strategy. (**B,D,F,H,J**) Monkey data in teal. More uncertain blocks had significantly smaller H(Str), ERDS, and EODS, indicating a more consistent strategy. MIOS was also significantly smaller, indicating a decreased link between previous choice option and strategy.

**Figure S13.**
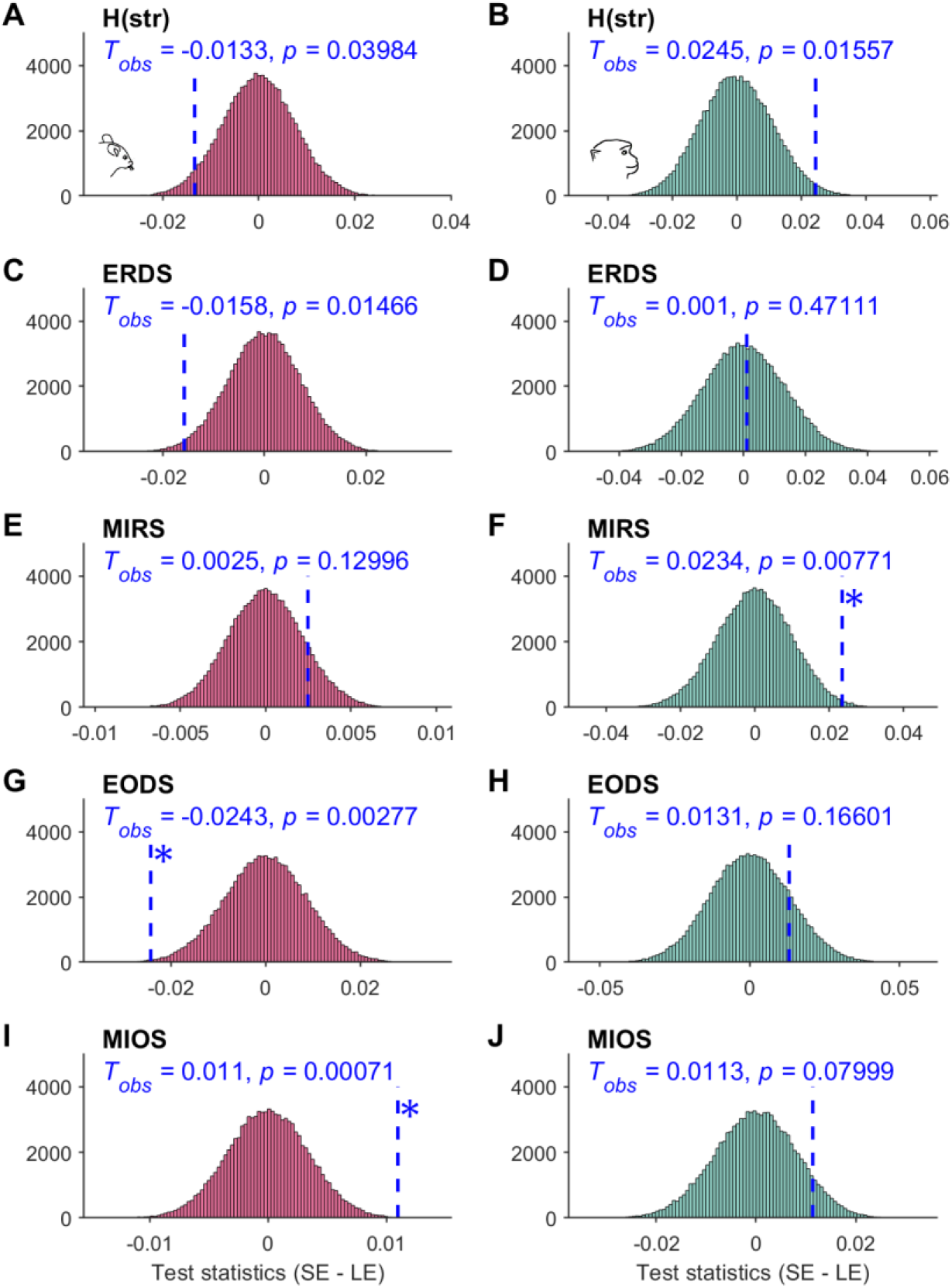
Effects of block switch expectation on mouse and monkey behavior shown by permutation test. Plots show distributions of test statistics from *N* = 100,000 sampled permutations in mice (column 1) and monkeys (column 2), under the null hypothesis that SE and LE blocks have the same entropy measures. Vertical dotted lines in blue indicate the actual observed test statistics (T_obs_), which is the difference between a given entropy metric for the two block types. P-values indicate the probability of observing test statistics as extreme as the observed value (one-tailed). Asterisks indicate *p* < 0.01. For each permutation, we randomly assigned blocks into two groups with sample size *n_1_* or *n_2_*, corresponding to the total number of blocks in SE or LE blocks, respectively. A single entropy measure is computed for each type of block by concatenating all trial vectors from the sampled permutation. Test statistic for each sample is then computed by taking the difference in metrics between the first and the second group. (**A,C,E,G,I**) Mouse data in red. LE blocks had significantly larger EODS (*Tobs* = −0.0243, *p* = .00241) and smaller MIOS (*T_obs_* = 0.011, *p* = .00061), indicating a change in option-dependent strategy. (**B,D,F,H,J**) Monkey data in teal. LE blocks had significantly smaller MIRS (*T_obs_* = 0.0131, *p* = .00762), indicating a change in reward-dependent strategy.

